# A Biopolymer Laminarin Elicits Antioxidant Defense in Different Cultivars of *Solanum Lycopersicum* Against Early Blight Disease Caused by *Alternaria Solani*

**DOI:** 10.1101/2025.10.05.680597

**Authors:** Govindan Muthukumar, Jeyapandi Mohana Prasad, Chinnasamy Arulvasu, Nallamuthu Godhantaraman, Mehanathan Muthamilarasan, Nagarathnam Radhakrishnan

## Abstract

The study evaluates the potential of laminarin (LaM), a biopolymer, in inducing antioxidative defense in tomato against early blight disease caused by *Alternaria solani*. Histological analysis of *A. solani* infection in susceptible (PKM-1) and tolerant (Arka Rakshak) cultivars of tomato showed increased pathogen colonization in PKM-1 by day 8 compared to Arka Rakshak. Fluorescent microscopic analysis with a chitin-specific dye, Calcofluor white, demonstrated reduced *A. solani* colonization in PKM-1 pretreated with LaM (0.1%), followed by *A. solani* infection on par with Arka Rakshak. A significant increase in the accumulation of hydrogen peroxide was observed in LaM-pretreated leaves (0.050%, 0.075% and 0.1%) at 24- and 48-hours post-infection in both cultivars. A quantitative increase in guaiacol peroxidase (GPX) activity was recorded in LaM (0.025%), followed by A. solani infection leaves at 36 hrs in Arka Rakshak, whereas LaM (0.010%) followed by *A. solani* infection recorded maximum GPX activity at 24 hrs in PKM-1. Induction of new GPX isoform was evident in LaM (0.075% and 0.1%) pretreated, and LaM pretreated, followed by *A. solani* infected leaves of both cultivars at 24 hrs. There was a significant reduction in disease severity in tomato leaves pretreated with LaM (0.1%) and infected with *A. solani*. The results suggest that LaM could be a potential elicitor of antioxidant defense in tomato against *A. solani* infection, which can further be harnessed to develop sustainable strategies for managing early blight disease while reducing dependence on chemical fungicides.

## 1. INTRODUCTION

*Solanum lycopersicum* L. (tomato), a member of the Solanaceae family, holds significant global importance as a widely cultivated vegetable crop due to its remarkable nutritional value. Tomatoes are a rich source of antioxidants, including vitamins C and E, organic acids such as malate and citrate, sugars like sucrose and hexoses, and bioactive compounds such as carotenoids and phenolics (Li *et al*., 2018). Despite its agricultural and nutritional significance, tomato production is severely impacted by various diseases, posing challenges to yield and quality. Among the major diseases affecting tomato crops, early blight, caused by *Alternaria solani*, is one of the most destructive, leading to substantial reductions in both fruit quality and yield (Rashmi *et al*., 2012). Plants have evolved robust antioxidant defense systems to mitigate the effects of reactive oxygen species (ROS), which are generated during biotic and abiotic stress responses. This antioxidant network comprises enzymatic components such as superoxide dismutase (SOD), catalase (CAT), and guaiacol peroxidase (GPX), which scavenge excess ROS and protect cellular integrity (Mittler *et al*., 2004; Mani *et al*., 2021). SOD catalyzes the dismutation of superoxide radicals (O2·-) into hydrogen peroxide (H2O2), which is subsequently detoxified into water and oxygen by CAT and GPX (Zhao *et al*., 2016). Furthermore, plants employ specialized mechanisms, including vacuolar compartmentalization, to detoxify excess ROS, highlighting the multifaceted nature of their stress adaptation strategies (Das and Roychoudhury, 2014; Mani *et al*., 2021).

The activation of plant defense mechanisms is tightly regulated by phytohormones, including abscisic acid (ABA), jasmonic acid (JA), ethylene (ET), and salicylic acid (SA), which modulate signaling pathways and transcriptional responses (War *et al*., 2011). Among these, SA plays a pivotal role in orchestrating defense responses by regulating key antioxidant enzymes such as SOD and peroxidases (War *et al*., 2011). Additionally, transcription factors (TFs) act as central mediators in transcriptional reprogramming, driving the synthesis of defense-related proteins and non-protein compounds (Jones and Dangl, 2006; Javed *et al*., 2020). The increasing reliance on synthetic fungicides for disease management raises concerns about environmental sustainability and food safety (Koley *et al*., 2015). Consequently, there is a growing interest in the development of eco-friendly alternatives, such as elicitor-based strategies, to enhance plant defense mechanisms.

Elicitors are molecules that trigger plant immune responses and are derived from various sources, including microorganisms, plants, and algae (Wiesel *et al*., 2014). Among these, algal-derived polysaccharides have emerged as promising bio-stimulants. Laminarin, a β-glucan polysaccharide derived from brown algae, has demonstrated the ability to enhance plant resistance by modulating hormonal signalling pathways and activating *defense*-related responses (Shukla *et al*., 2021). In *Camellia sinensis*, laminarin influences salicylic acid and MAPK signalling, contributing to enhanced resistance against pests (Xin *et al*., 2019). Similarly, in *Arabidopsis thaliana*, laminarin stabilizes chloroplast function under abiotic stress conditions, underscoring its multifaceted role in plant *defense* and development (Wu *et al*., 2016). This study aims to evaluate the efficacy of laminarin as a biopolymer elicitor in inducing defense responses against early blight disease in tomato plants caused by *A. solani*. By elucidating the biochemical and molecular mechanisms involved, this research seeks to contribute to the development of sustainable disease management strategies for tomato cultivation.

## 2. MATERIALS AND METHODS

### 2.1 Plant materials: Seed source and seed variety

Seeds of the early blight-susceptible tomato cultivar PKM-1 (*Solanum lycopersicum*) were procured from the Horticultural College and Research Institute (HRI-TNAU), Periyakulam, Theni District, Tamil Nadu, India. The early blight-tolerant cultivar, Arka Rakshak, was obtained from the Indian Institute of Horticultural Research (IIHR), Bengaluru, Karnataka, India. Plants were grown under controlled greenhouse conditions with a 16/8-hour (Day/Night) photoperiod at 25°C in pots containing a mixture of commercial manure and vermiculite (TAMIN, Chennai). The plants were maintained until the 4–5 leaf stage (4 weeks old) for subsequent elicitor treatments and *A. solani* infection (Mani and Nagarathnam, 2018).

### 2.2 Pathogenicity experiment

The pathogenicity test was conducted using *A. solani* fungal cultures on four-week-old healthy tomato plants cultivated in sterile soil under glasshouse conditions. *A. solani* was grown on corn meal agar (CMA) in darkness for 12–15 days. A spore suspension (2 × 10□ mL□^1^) was prepared and sprayed onto the leaves of experimental plants until runoff. Control plants were sprayed with sterile double-distilled water. To ensure infection, treated plants were incubated under polythene covers at 25°C, 80–100% relative humidity, and a 16/8-hour (Day/Night) photoperiod. Disease symptoms (necrotic spots) were monitored daily for 8 days, and Koch’s postulates were confirmed (Gao *et al*., 2017).

### 2.3 Visual examination of disease symptoms

Symptoms of early blight, characterized by necrotic spots, were visually assessed and documented in *A. solani*-infected tomato plants. Observations were made on control plants, *A. solani*-infected plants, and plants treated with laminarin alone or laminarin followed by *A. solani* infection at two-day intervals up to 8 days post-treatment.

### 2.4 Elicitor treatment

Commercially available laminarin (LaM) (YL76431, Biosynth® Carbosynth, United Kingdom) was used as the elicitor. Foliar applications of laminarin were performed on 30-day-old greenhouse-grown plants. Different concentrations of laminarin (0.010%, 0.025%, 0.050%, 0.075%, and 0.1% w/v) were prepared in sterile double-distilled water and sprayed until runoff on both adaxial and abaxial leaf surfaces. Control plants were sprayed with sterile double-distilled water. All treated plants were maintained in greenhouse conditions with a 16/8-hour (Day/Night) photoperiod at 25°C and left undisturbed for 48 hours before further experiments.

### 2.5 Histo-pathological studies of *A. solani* infected tomato leaves

#### 2.5.1 Light microscopy

After the appearance of the first visible symptoms of *A. solani* infected *S. lycopersicum* leaves (Day ‘0’), the leaves were collected at 2, 4, 6, and 8-day intervals for microscopic observations. The leaves were decolorized in acetic acid and ethanol (1:3) solution during overnight incubation at 25°C. Subsequently, the leaves were rinsed twice with autoclaved distilled water and stained using a 0.05% lactophenol cotton blue solution.

After staining, the leaves were mounted on a glass slide with 40% glycerol. Qualitative observations of *A. solani* infected tomato leaves, including hyphal distribution and penetration, were made using a light microscope (Thinakaran *et al*., 2020).

#### 2.5.2 Fluorescence microscopy

To study the penetration of pathogens on tomato leaves showing disease symptoms after infection with *A. solani*, the experimental tomato leaves were stained with calcofluor white stain (Sigma Aldrich, USA). The leaves were placed on a clean glass slide on which a drop of calcofluor white stain (Calcofluor white M2R 1g/L and Evans blue 0.5 g/L) and Propidium iodide (1mg/mL) were added as a counter stain. The slides were examined under a fluorescence microscope (Nikon Corporation, Japan) using a DAPI filter (EX 362-396, DM 415 and BA 432-482) and then TRITC (EX 532-554, DM 565 and BA 574-626) at excitation/emission wavelengths.

### 2.6 Accumulation of ROS

#### 2.6.1 *In-situ* Histo-chemical staining of Hydrogen peroxide (H_2_O_2_) accumulation

A freshly prepared 3,3’-Diaminobenzidine (DAB) staining solution was used to visually analyse the hydrogen peroxide of experimental tomato leaves (H_2_O_2_) (Daudi and Brien, 2012). The experimental leaves (three leaves each) were soaked in DAB solution for 4-5 hrs in a shaker (80 rpm). Then, the leaves were submerged in a bleaching solution (Ethanol, acetic acid and glycerol (3:1:1) for 30 min. The leaves were then decoloured by boiling the samples with a bleaching solution. The resultant brown colour formation due to the interaction of DAB with hydrogen peroxide was observed and photographed.

#### 2.6.2 *In-situ* Histo-chemical detection of superoxide anion (O2•−) accumulation

A freshly prepared Nitro Blue Tetrazolium (NBT) staining solution was used to visually analyse the superoxide anion radical in the experimental leaves of tomato (Kumar *et al*., 2014). All the experimental leaves were stained with NBT incubated at room temperature and added 0.05% NBT solution to submerge the leaves. The plate was placed on a typical laboratory shaker for 4 to 5 hrs at 80 to 100 rpm. The reaction was stopped by exposing the leaves to a hot water bath to which 95% ethanol was added to remove excess colours. When superoxide anions react with NBT in leaves, the colour fades. After boiling, a dark blue colour appears. The visible colour change was photographed while illuminated in white light.

### 2.7 Antioxidant enzyme assays

#### 2.7.1 Guaiacol Peroxidase (GPX)

The activity of Guaiacol Peroxidase (GPX) in all of the experimental leaves was tested using the Volk and Feierabend (1989) method. In a reaction mixture of 3 mL, containing 50μL of enzyme extract, 2.93mL of sodium phosphate buffer, 9.8μL of guaiacol (30mM), and 2μL of H2O2 (6.5mM). The GPX absorbance was measured at 470 nm. Analysis of GPX isoenzymes was performed by 7% Native PAGE. Following a 30-min immersion in 0.018M guaiacol, the gel was washed with deionized water and then exposed to 1% (v/v) glacial acetic acid and 0.015% H_2_O_2_ until dark brown bands formed (Mani *et al*., 2021).

#### 2.7.2 Polyphenol oxidase (PPO)

The PPO activity of all experimental plant leaves was measured using the method described by Arnnok *et al*., (2010). PPO isoform exploration using native-PAGE (7%) gel stained with 100mM catechol for 30 minutes (gel rocker). The gels were then washed in distilled water, then stored at 4°C with 5% (v/v) acetic acid.

#### 2.7.3 Superoxide Dismutase (SOD)

SOD activity was assessed using the Beauchamp and Fridovich (1971) method. In a reaction volume of 3mL contains 70μl of supernatant leaf extract, 2μL of EDTA (0.05mM), 114μL of methionine (13mM), 23μL of NBT (75mM), and 9μL of riboflavin (20μM). SOD activity was measured at 560 nm after a 10-minute incubation period. Analysis of SOD isoenzymes gel was immersed with Tris-HCl buffer, Na_2_ EDTA and NBT, riboflavin 0.1M in the dark at 25°C for 30 min. The gel was further illuminated (white light) in 15 min (Dey *et al*., 2019).

#### 2.7.4 Catalase (CAT)

All of the experimental tomato leaves were assayed CAT activity by Volk and Feierabend (1989) method. The reaction mixture contains 60μL of enzyme extract, 5μL of H2O2 (14mM), and 2.93mL of potassium phosphate buffer (0.1M; pH 7.0). CAT activity was measured by Δ240 nm.

#### 2.7.5 Phenylalanine ammonia-lyase (PAL)

Phenylalanine ammonia-lyase activity was determined using the Lavania *et al*., (2006) method. 100μL of leaf extract was combined with 900μL of 0.5M Tris-HCl buffer and 6μM L-phenylalanine, then water bathed at 37°C for 70 min.

### 2.8 Disease scoring

Disease scoring was evaluated by observing the development of disease symptoms in the four week old tomato plants in all the experimental groups such as control, *A. solani* infected and laminarin pretreated and followed by *A. solani* infected groups. Scale of disease scoring was done using Disease Scoring Scale of 0 Scale: No symptoms on the leaf, 1 Scale: ≤ 25% leaf area infected and covered by spots, 2 Scale: 26 ≤ 50% leaf area infected and covered by spot, branches, 3 Scale: 51 ≤ 75%leaf area infected and covered by spot and 4 Scale: 76 ≤ 100% leaf area infected and covered by spot.

### 2.9 Statistical analysis

Experimental results obtained in the study on *A. solani* infection, laminarin pretreated alone, and laminarin pretreatment followed by *A. solani* infection treatment were subjected to apt statistical analyses. The data were subjected to a two-way analysis of variance (ANOVA) followed by Tukey’s multiple comparisons tests. The statistical analysis was performed by GraphPad Prism version 9.0 (GraphPad Software, San Diego, CA, USA) (Irulappan *et al*., 2022).

## 3. RESULTS

### 3.1 Koch’s postulates and morphological identification of *Alternaria solani*

The initial symptoms of early blight disease in tomato plants were observed as small, brown necrotic spots with water-soaked lesions on older leaves. These lesions progressively spread upward from the bottom to the top of the plant. By day 7, the necrotic spots had enlarged, assuming an angular or oval shape with a surrounding thin, yellow, chlorotic halo. Over time, the spots became more prominent, developing concentric rings that created a distinctive bullseye pattern. Adjacent spots coalesced, forming larger, irregular patches. The re-isolation and culturing of *A. solani* from the re-infected leaves confirmed its morphological identity, which matched the original fungal culture. The morphological characteristics were consistent with the descriptions outlined in *Alternaria: An Identification Manual* (Simmons, 2007). These findings confirmed the pathogenicity of *A. solani* and supported Koch’s postulates (Fig. 1).

**Fig. 1.**
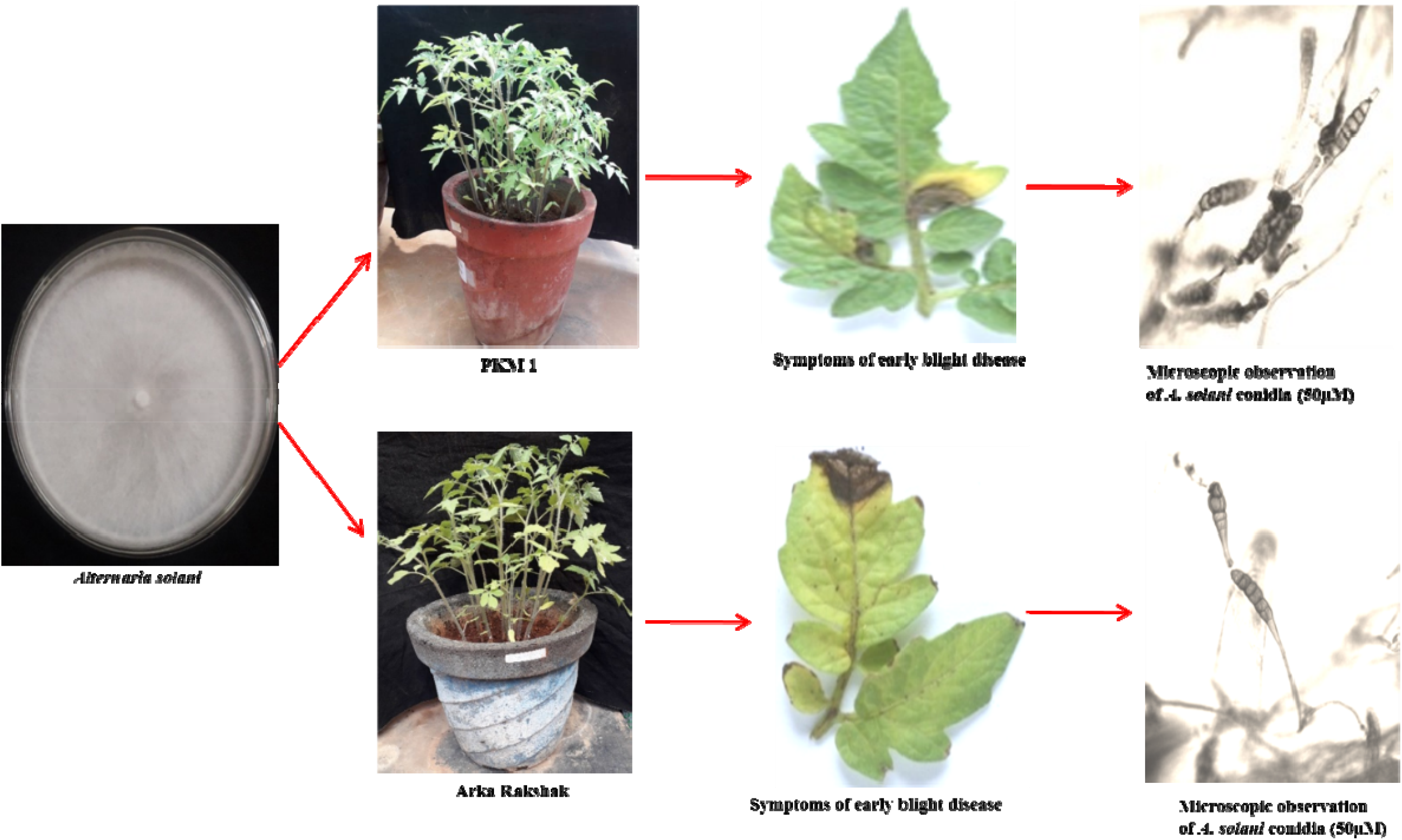
Koch’s postulates test of pathogenicity in *S. lycopersicum* infected with *A. solani*.

### 3.2 Symptoms of *A. solani* infection on *S. lycopersicum*

On day 7, early symptoms of blight disease were observed in both tolerant and susceptible cultivars of *S. lycopersicum*. These symptoms manifested as tiny, angular, brown necrotic patches with concentric rings surrounded by yellow halos. In contrast, no symptoms were observed in the control group or in plants pretreated with laminarin. Day 0 was marked as the time when the first symptoms appeared. In the susceptible cultivar PKM-1, a dose- and time-dependent reduction in disease symptoms was evident in leaves pretreated with various concentrations of laminarin (0.010%, 0.025%, 0.050%, 0.075%) by day 8. The most pronounced delay in symptom onset occurred in plants treated with 0.1% laminarin, where necrotic lesions appeared only by day 6. In the tolerant cultivar Arka Rakshak, a marked reduction in symptom progression was observed in plants pretreated with laminarin, indicating enhanced resistance to *A. solani* infection (Fig. 2A & 2B).

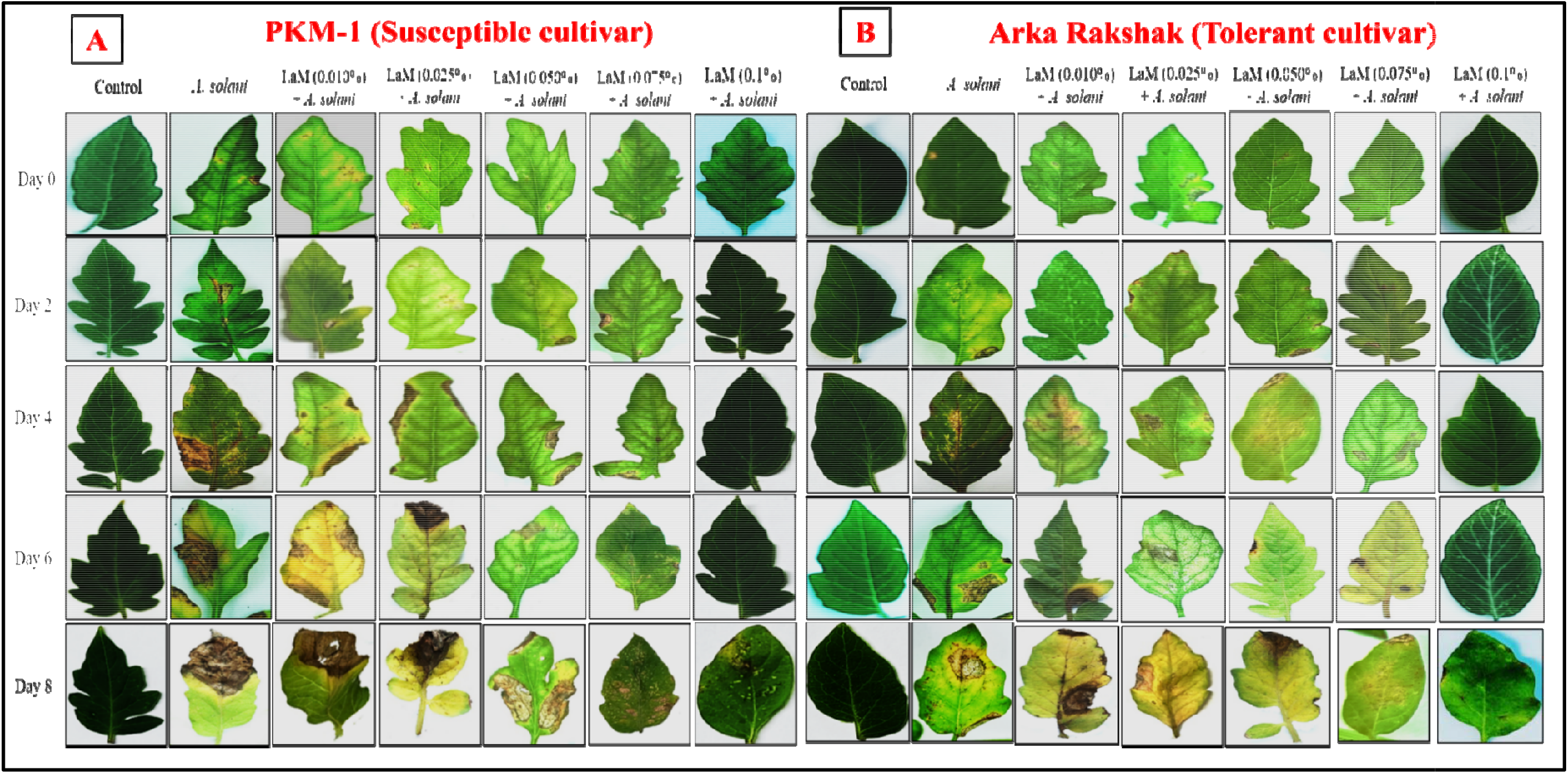
**Fig. 2A**. Typical symptoms of early blight disease *S. lycopersicum* (PKM-1, susceptible cultivar) leaves pretreated with various concentrations of laminarin followed by *A. solani* infected samples collected after the first visible symptoms appeared. **Fig.2B**. Typical symptoms of early blight disease *S. lycopersicum* (Arka Rakshak, tolerant cultivar) leaves pretreated with various concentrations of laminarin followed by *A. solani* infected samples collected after the first visible symptoms appeared.

### 3.3 Light microscopy observation

Lactophenol cotton blue staining was utilized to study the development of *A. solani* within the leaves of both susceptible and tolerant tomato cultivars. In susceptible cultivar PKM-1, fungal hyphal amplification was evident from day 2 through day 8, displaying progressive colonization and ramification within the leaf tissues. However, laminarin pretreatment at 0.1% significantly inhibited fungal proliferation, with fungal structures only becoming visible on day 8. In contrast, the tolerant cultivar Arka Rakshak exhibited limited fungal growth even in the absence of laminarin pretreatment, suggesting an innate resistance mechanism. In laminarin-treated leaves of the tolerant cultivar (0.1%), fungal growth was completely suppressed, and no visible signs of infection were observed throughout the experimental period. These findings highlight a dose- and time-dependent suppression of fungal growth in laminarin-treated susceptible cultivars and enhanced resistance in the tolerant cultivar (Fig. 3A & 3B).

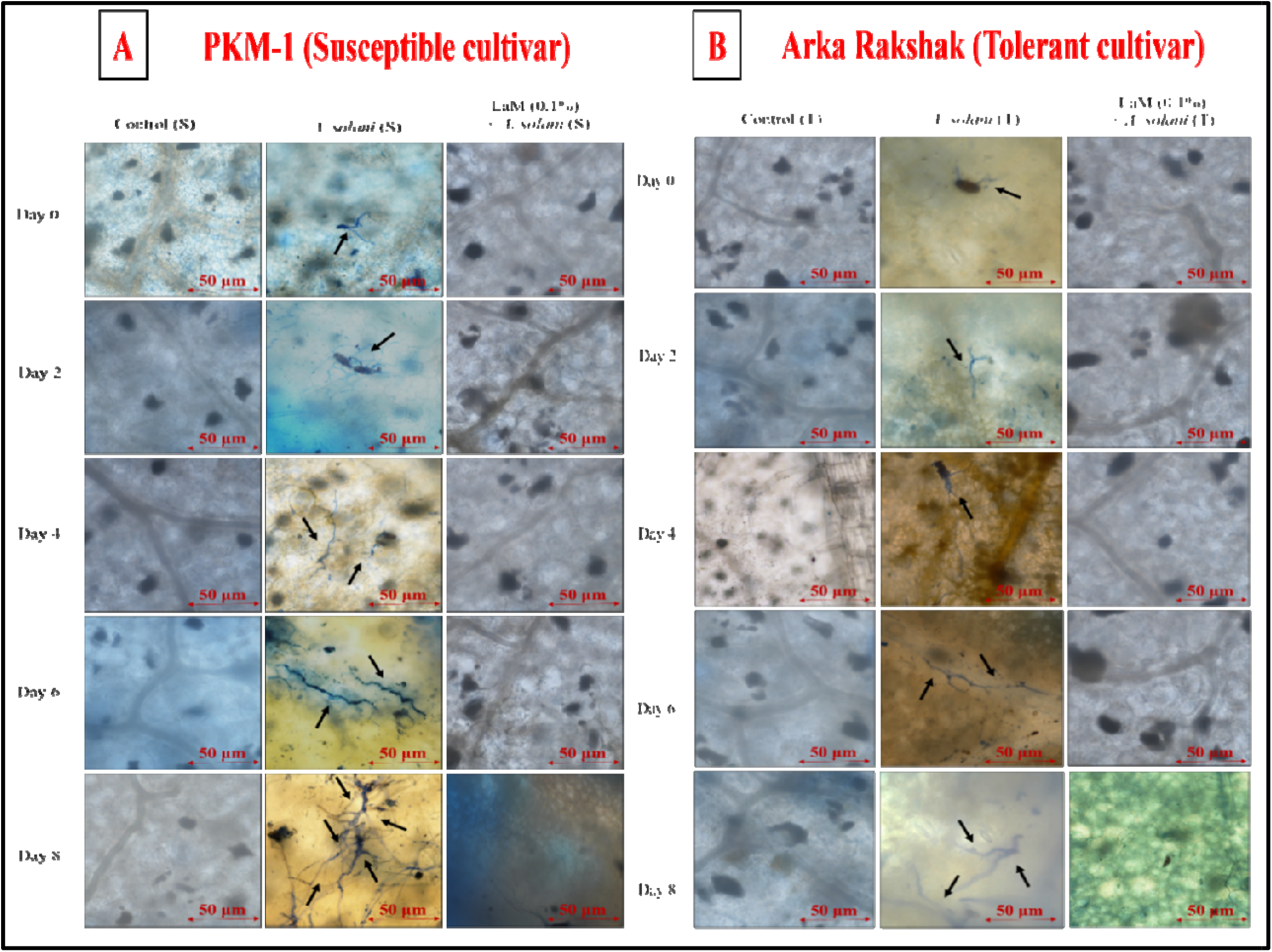
**Fig. 3A**. Microscopic analysis of *A. solani* infection patterns on *S. lycopersicum* (PKM-1, susceptible cultivar) pretreated with laminarin and sample collected after the first visible symptoms appeared. **Fig.3B**. Microscopic analysis of *A. solani* infection patterns on *S. lycopersicum* (Arka Rakshak, tolerant cultivar) pretreated with laminarin and sample collected after the first visible symptoms appeared.

### 3.4 Fluorescence microscopy observation

To examine fungal penetration and colonization, tomato leaves were stained with calcofluor white and propidium iodide and observed under fluorescence microscopy. In susceptible cultivar PKM-1, *A. solani* displayed significant penetration and colonization by day 7, with a progressive increase in fungal structures from day 2 to day 8. Laminarin pretreatment significantly reduced fungal penetration, with complete suppression of colonization in tolerant cultivars pretreated with 0.1% laminarin. In the tolerant cultivar Arka Rakshak, fungal colonization was notably reduced even without laminarin pretreatment, while pretreatment with 0.1% laminarin resulted in the absence of fungal structures throughout the experimental period. These results demonstrate that laminarin effectively mitigates fungal invasion and enhances resistance in both susceptible and tolerant cultivars (Fig. 4A & 4B, S1–S3).

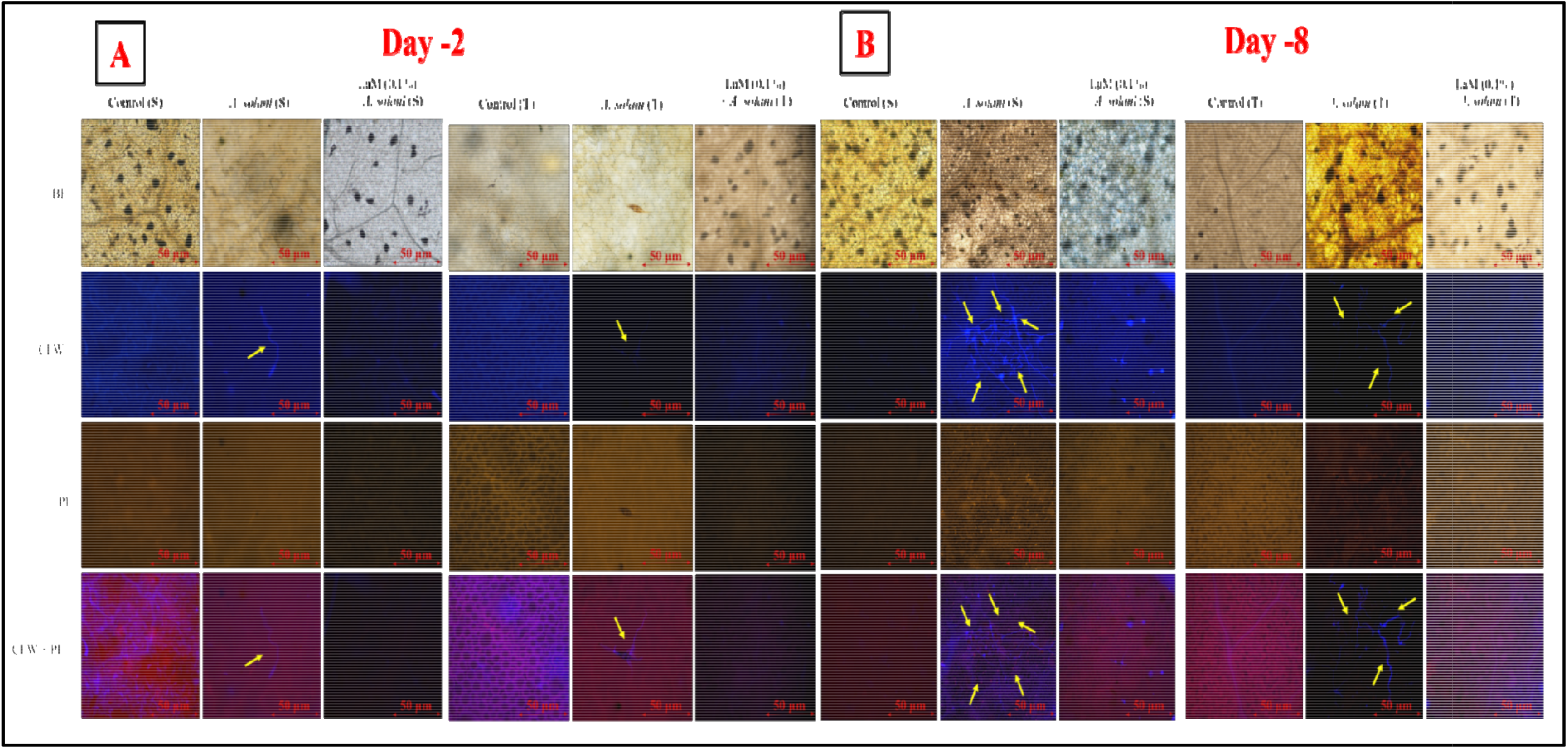
**Fig. 4A**. Fluorescence microscopic analysis of *A. solani* infection patterns on *S. lycopersicum* (PKM-1, susceptible cultivar) and (Arka Rakshak, tolerant cultivar) pretreated with laminarin and followed by *A. solani*. Samples collected after the first visible symptoms appeared S. susceptible cultivar and T. tolerant cultivar. at day 2. **Fig.4B**. Fluorescence microscopic analysis of *A. solani* infection patterns on *S. lycopersicum* (PKM-1, susceptible cultivar) and (Arka Rakshak, tolerant cultivar) pretreated with laminarin and followed by *A. solani*. Samples collected after the first visible symptoms appeared S. susceptible cultivar and T. tolerant cultivar. at day 8.

### 3.5 *In-Situ* histochemical localization of hydrogen peroxide (H□O□)

The accumulation of hydrogen peroxide (H□O□) was studied using in-situ histochemical staining with DAB. A time-dependent increase in H□O□ accumulation was observed in tomato leaves pretreated with laminarin and/or infected with *A. solani*. The tolerant cultivar exhibited a more substantial and earlier accumulation of H□O□ compared to the susceptible cultivar. Laminarin pretreatment induced a dose-dependent enhancement of H□O□ levels, with the highest accumulation recorded in plants treated with 0.1% laminarin. This increase was observed across both susceptible and tolerant cultivars, but the tolerant cultivar showed significantly higher H□O□ levels at all time points. Minimal effects were observed in plants treated with the lowest concentration of laminarin (0.010%), indicating that higher concentrations are required for significant oxidative responses (Fig. 5A & 5B).

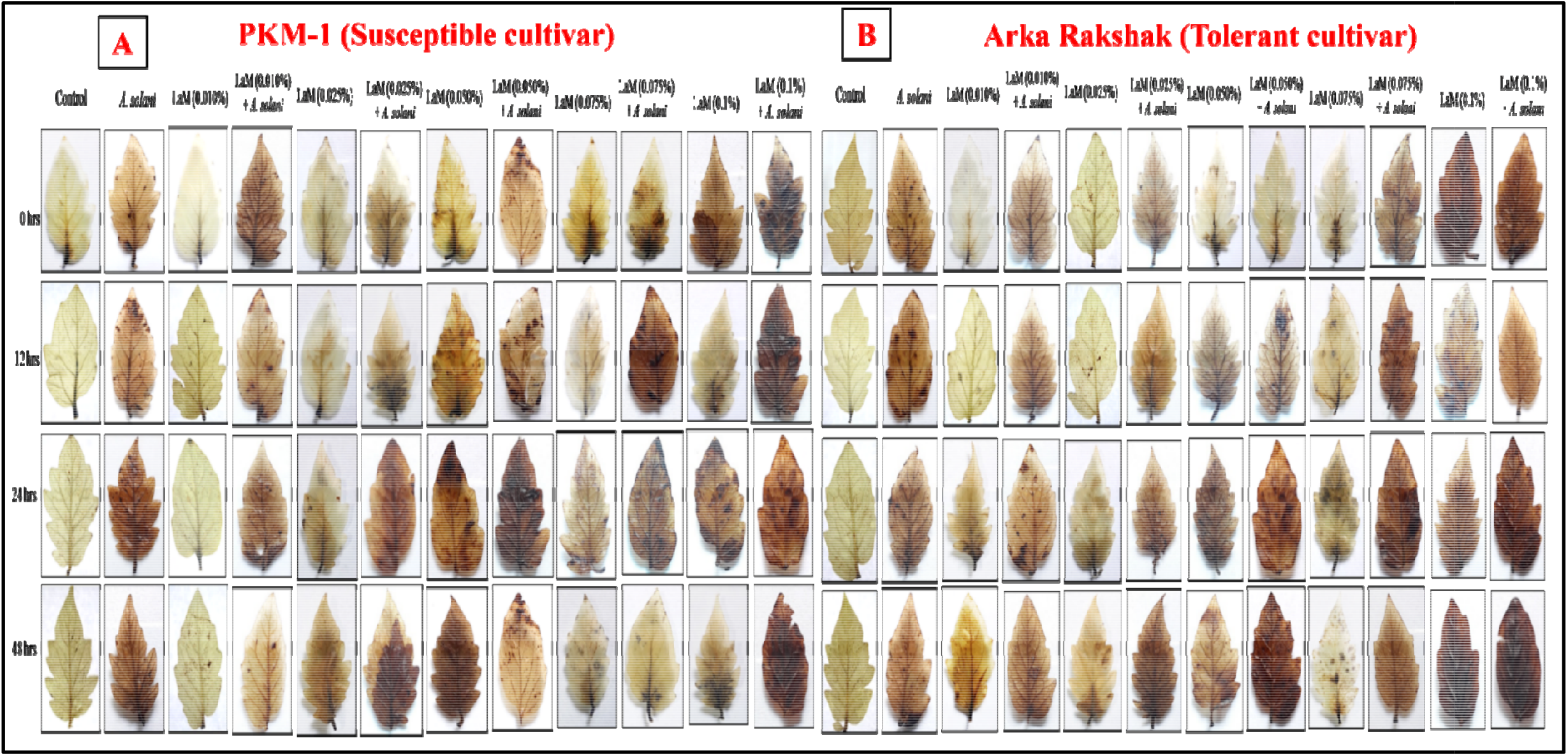
**Fig. 5A**. *In-situ* histochemical localization of hydrogen peroxide (H_2_O_2_) accumulation on *S. lycopersicum* (PKM-1, susceptible cultivar) leaves pretreated with different concentrations of laminarin and samples collected at 0 hrs, 12 hrs, 24 hrs and 48 hr. **Fig.5B**. *In-situ* histochemical localization of hydrogen peroxide (H_2_O_2_) accumulation on *S. lycopersicum* (Arka Rakshak, tolerant cultivar) leaves pretreated with different concentrations of laminarin and samples collected at 0 hrs, 12 hrs, 24 hrs and 48 hr.

### 3.6 *In-Situ* histochemical localization of superoxide anion (O□ · □)

The accumulation of superoxide anion (O□·□) was visualized using NBT staining. A dose- and time-dependent increase in O□·□ accumulation was evident in laminarin-pretreated leaves. Tomato leaves infected with *A. solani* exhibited significant O□·□ accumulation in both cultivars, with the susceptible cultivar displaying higher levels. However, laminarin pretreatment markedly enhanced O□·□ levels in both cultivars, with the highest accumulation observed in leaves treated with 0.1% laminarin. In the tolerant cultivar, this response was sustained over time, indicating an effective defense mechanism against *A. solani* infection. Minimal effects were observed in plants treated with 0.010% laminarin, confirming the importance of optimal laminarin concentrations for significant defense activation (Fig. S4A & S4B).

### 3.7 Effect of laminarin on antioxidant enzyme activities

#### 3.7.1 Guaiacol peroxidase (GPX) activity

Laminarin pretreatment significantly enhanced GPX activity in both susceptible and tolerant cultivars, with a pronounced dose- and time-dependent increase observed at a 0.1% concentration. The tolerant cultivar showed a greater rise in GPX activity compared to the susceptible cultivar. New GPX isoforms were induced in both cultivars, particularly at laminarin concentrations of 0.075% and 0.1%, following *A. solani* infection (Fig. 6A & 6B, Fig. 7, S5).

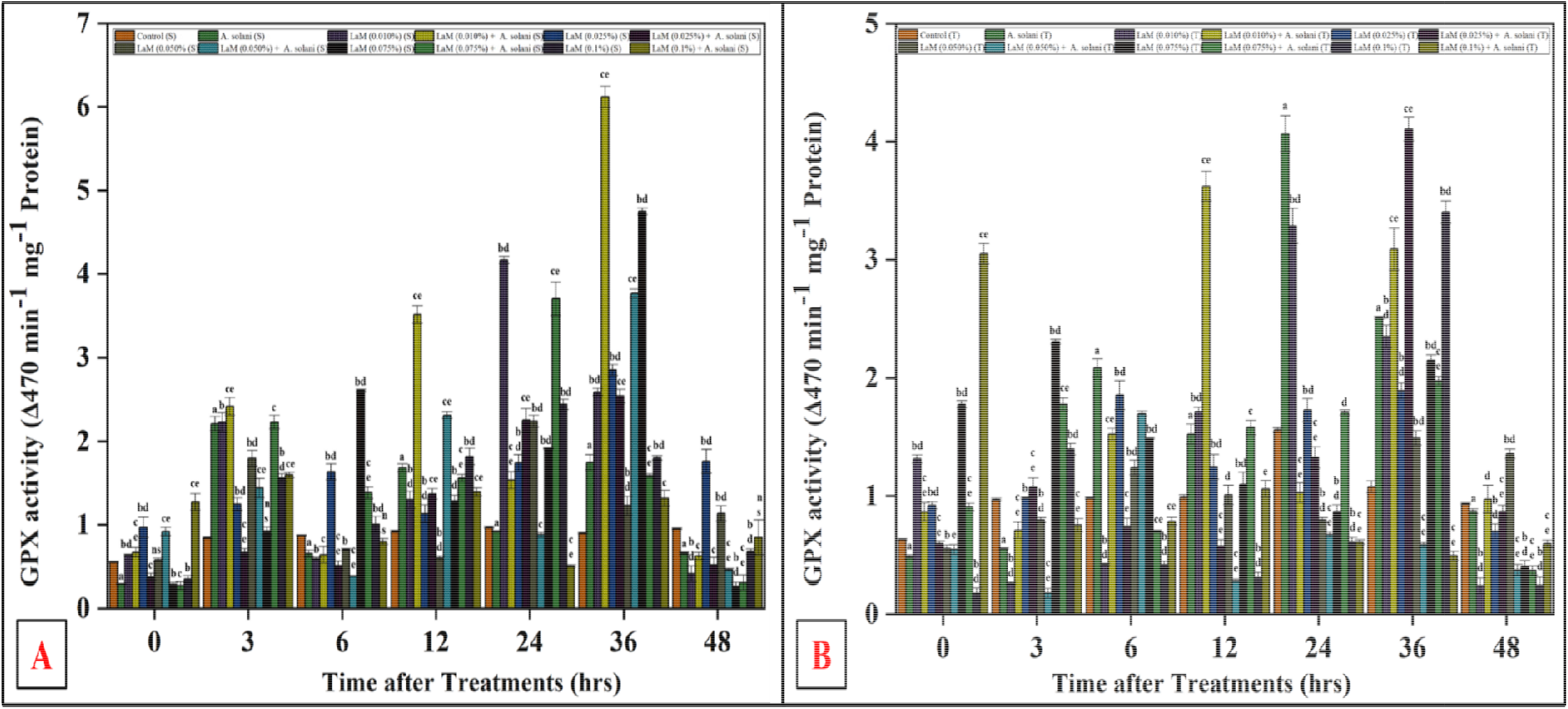
**Fig. 6A**. Guaiacol peroxidase (GPX) activity of *S. lycopersicum* (PKM-1, susceptible cultivar) leaves pretreated with different concentrations of laminarin and infected with *A. solani*. **Fig.6B**. Guaiacol peroxidase (GPX) activity of *S. lycopersicum* (Arka Rakshak, tolerant cultivar) leaves pretreated with different concentrations of laminarin and infected with *A. solani*.

**Fig. 7.**
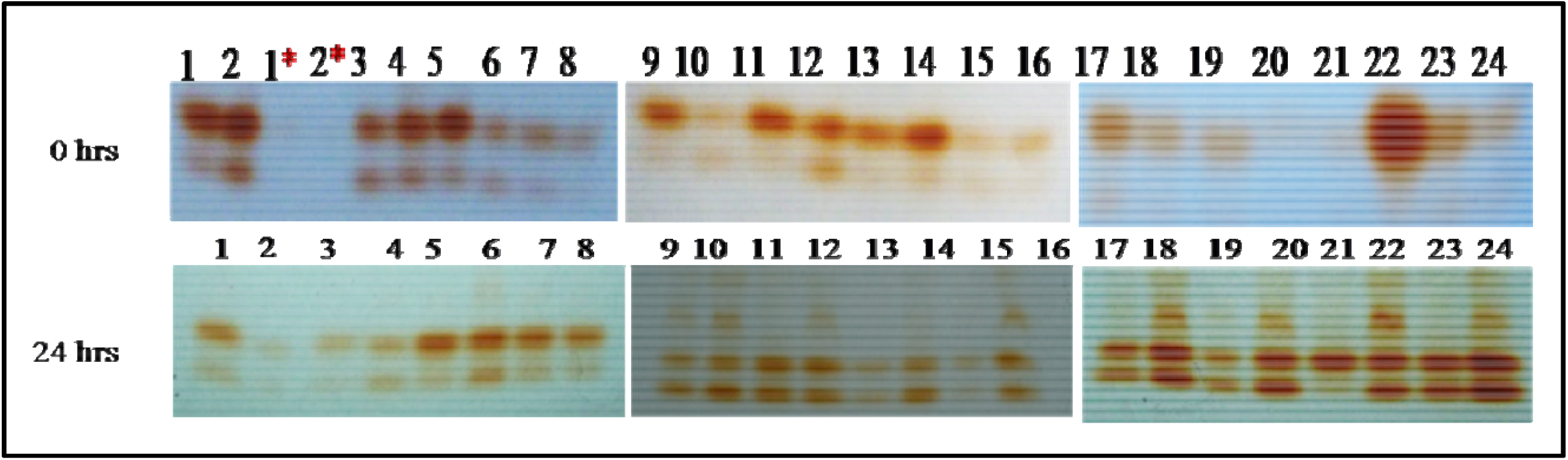
Native-PAGE analysis of Guaiacol peroxidase (GPX) isozymes expressions of *S. lycopersicum* leaves pretreated with different concentrations of laminarin and infected with *A. solani* 1. Control (S), 2. Control (T), 1*. Control (S) (GPX inhibitor, Sodium azide (15%)), 2*. Control (T) (GPX inhibitor, Sodium azide (15%)), 3. *A. solani* infected (S), 4. *A. solani* infected (T), 5. Laminarin (0.010%) (S), 6. Laminarin (0.010%) (T), 7. Laminarin (0.010%) + *A. solani* infected (S), 8. Laminarin (0.010%) + *A. solani* infected (T), 9. Laminarin (0.025%) (S), 10. Laminarin (0.025%) (T), 11. Laminarin (0.025%) + *A. solani* infected (S), 12. Laminarin (0.025%) + *A. solani* infected (T), 13. Laminarin (0.050%) (S), 14. Laminarin (0.050%) (T), 15. Laminarin (0.050%) + *A. solani* infected (S), 16. Laminarin (0.050%) + *A. solani* infected (T), 17. Laminarin (0.075%) (S), 18. Laminarin (0.075%) (T), 19. Laminarin (0.075%) + *A. solani* infected (S), 20. Laminarin (0.075%) + *A. solani* infected (T), 21. Laminarin (0.1%) (S), 22. Laminarin (0.1%) (T), 23. Laminarin (0.1%) + *A. solani* infected (S), 24. Laminarin (0.1%) + *A. solani* infected (T). S - susceptible cultivar and T - tolerant cultivar. 22.

#### 3.7.2 Polyphenol oxidase (PPO) activity

PPO activity increased significantly in laminarin-pretreated leaves, with the highest activity observed at 0.1% concentration. This increase was time- and dose-dependent in both cultivars, with the tolerant cultivar exhibiting a more robust response. New PPO isoforms were detected in laminarin-treated leaves followed by *A. solani* infection, indicating an enhanced enzymatic defense response (Fig. S6A & S6B, Fig. S7).

#### 3.7.3 Superoxide dismutase (SOD) activity

SOD activity increased significantly in both cultivars following laminarin pretreatment. The tolerant cultivar showed consistently higher SOD activity compared to the susceptible cultivar. No changes in SOD isoenzyme patterns were observed, suggesting that laminarin influences existing enzymatic activity rather than inducing new isoforms (Fig. 8A & 8B, Fig. S8).

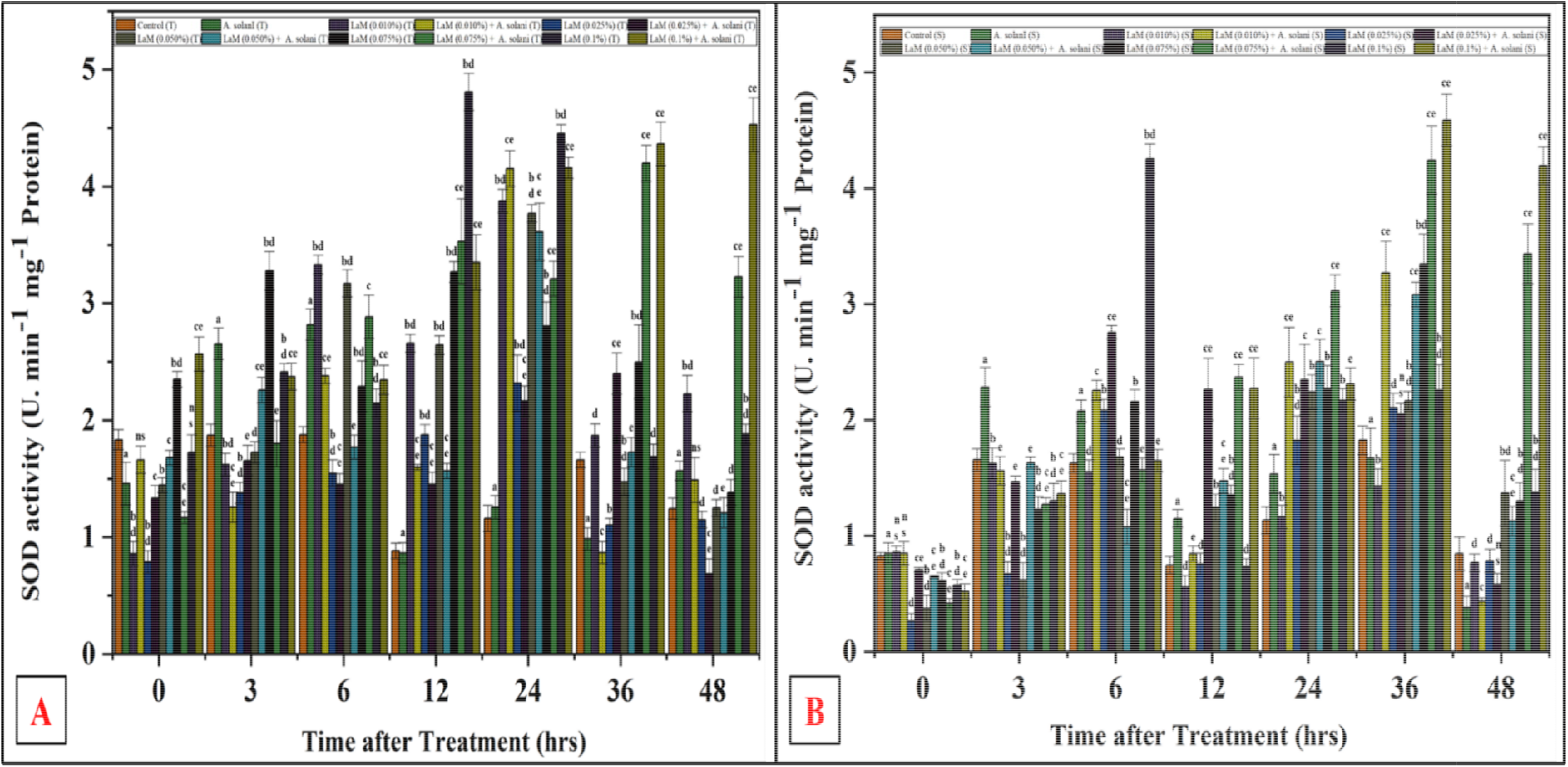
**Fig. 8A**. Superoxide Dismutase activity (SOD) activity of *S. lycopersicum* (PKM-1, susceptible cultivar) leaves pretreated with concentrations of laminarin and infected with *A. solani*. different **Fig.8B**. Superoxide Dismutase activity (SOD) activity of *S. lycopersicum* (Arka Rakshak, tolerant cultivar) leaves pretreated with different concentrations of laminarin and infected with *A. solani*.

#### 3.7.4 Catalase (CAT) activity

Catalase activity exhibited a dose- and time-dependent increase in laminarin-treated leaves, with a maximum response at 0.1% concentration. The tolerant cultivar consistently demonstrated higher CAT activity compared to the susceptible cultivar across all treatments (Fig. 9A & 9B).

**Fig. 9A.**
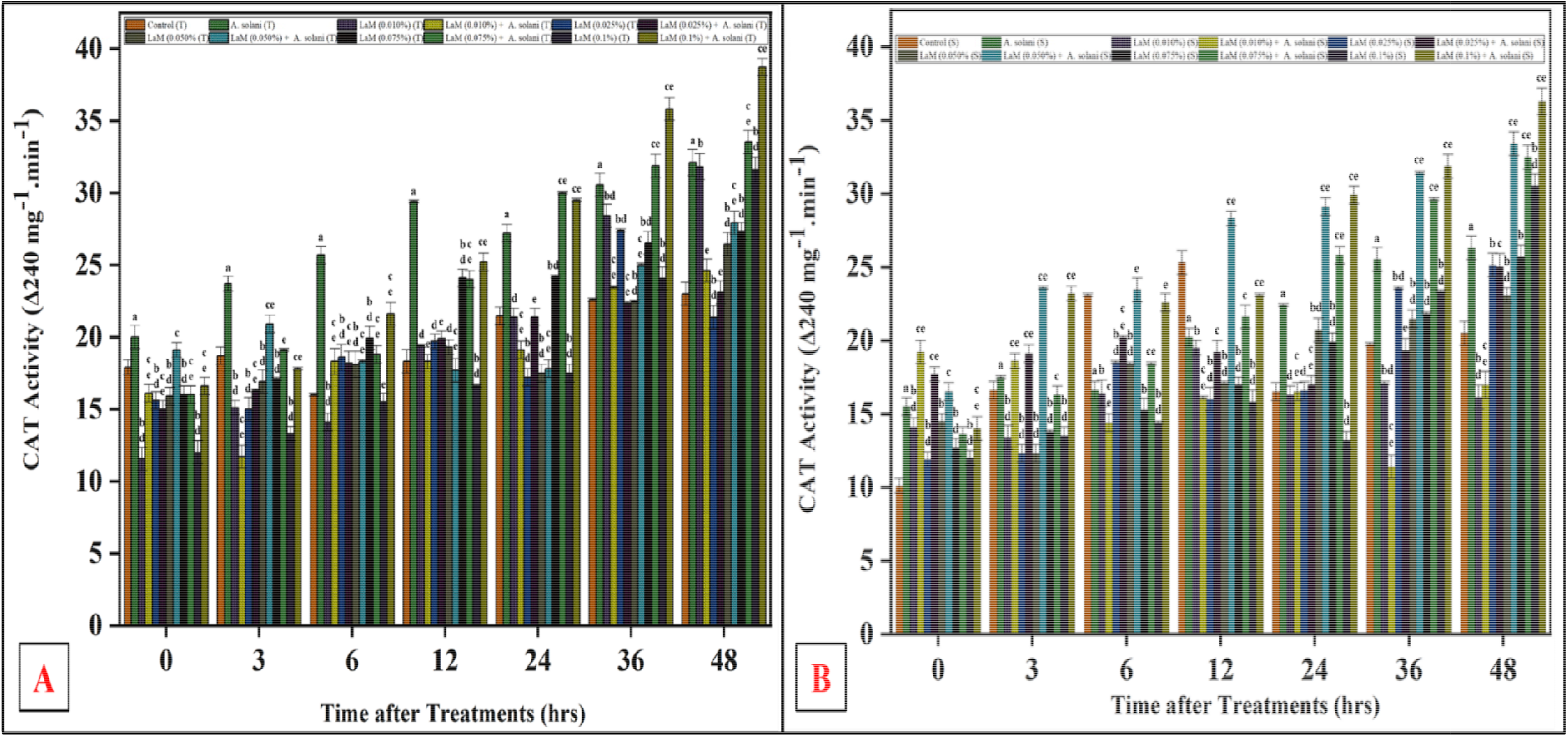
**Fig. 9A**. Catalase (CAT) activity of *S. lycopersicum* (PKM-1, susceptible cultivar) leaves pretreated with different concentrations of laminarin and infected with *A. solani*. **Fig.9B**. Catalase (CAT) activity of *S. lycopersicum* (Arka Rakshak, tolerant cultivar) leaves pretreated with different concentrations of laminarin and infected with *A. solani*.

#### 3.7.5 Phenylalanine ammonia-lyase (PAL) activity

PAL activity showed a significant increase in laminarin-pretreated leaves, with a pronounced time- and dose-dependent response. The tolerant cultivar exhibited higher PAL activity levels compared to the susceptible cultivar. The highest PAL activity was observed in plants treated with 0.1% laminarin (Fig. S9A & S9B).

### 3.9 Disease scoring

Disease severity was assessed using a scoring scale, and results indicated a progressive manifestation of symptoms in *A. solani*-infected tomato plants. By day 20, the susceptible cultivar PKM-1 recorded a disease severity score of 4 (76–100%), while the tolerant cultivar scored 3 (51–75%). Laminarin pretreatment at 0.1% resulted in a significant reduction in disease severity, with a final score of 1 (≤25%) in both cultivars. This highlights the efficacy of laminarin in mitigating disease progression (Fig. 10).

**Fig. 10.**
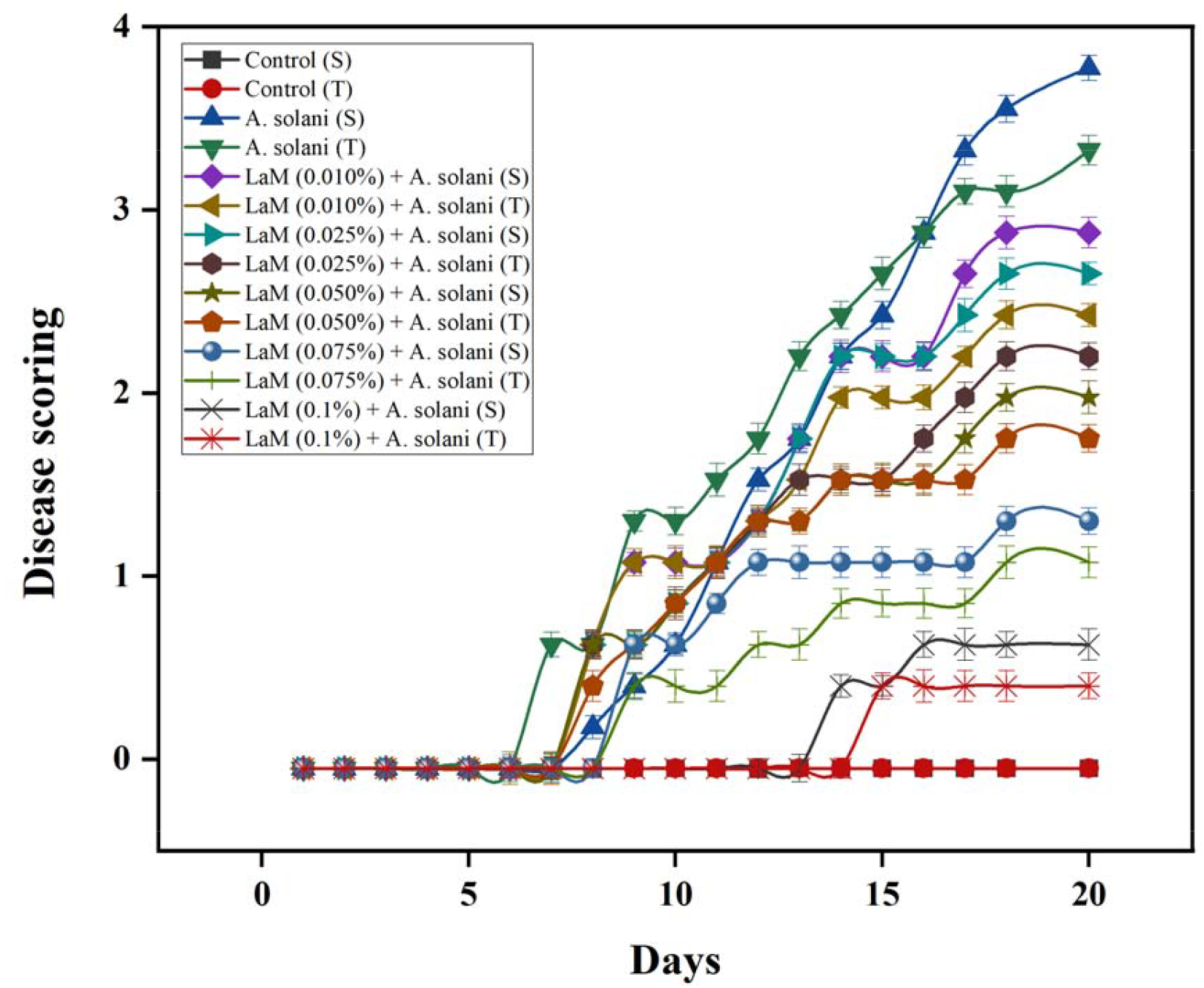
Disease Scoring of leaves of *S. lycopersicum* pretreated with laminarin and infected with *A. solani*. 1. Control (S), 2. Control (T), 3. *A. solani* infected (S), 4. *A. solani* infected (T), 5. Laminarin (0.010%) pretreated and followed by *solani* infected (S), 6. Laminarin (0.010%) pretreated and followed by *A. solani* infected (T), 7. Laminarin (0.025%) pretreated and followed by *A. solani* infected (S), 8. Laminarin (0.025%) pretreated and followed by *A. solani* infected (T), 9. Laminarin (0.050%) pretreated and followed by *A. solani* infected (S), 10. Laminarin (0.050%) pretreated and followed by *A. solani* infected (T), 11. Laminarin (0.075%) pretreated and followed by *A. solani* infected (S), 12. Laminarin (0.075%) pretreated and followed by *A. solani* infected (T), 13. Laminarin (0.1%) pretreated and followed by *A. solani* infected (S), 14. Laminarin (0.1%) was pretreated and followed by *A. solani* infected (T). S - susceptible cultivar and T - tolerant cultivar.

## 4. DISCUSSION

Algal polysaccharides have been extensively demonstrated as effective elicitors for inducing plant resistance against fungal infections and other pathogens (Dey *et al*., 2019). The current investigation revealed that tomato leaves pretreated with laminarin (0.1%) followed by infection with *A. solani* exhibited a significant reduction in disease progression by day 8 in both susceptible and tolerant cultivars. The pathogenicity tests confirmed that *A. solani* isolated from infected tomato leaves retained its pathogenicity, inducing characteristic disease symptoms on tomato plants as previously described (Mugao, 2023). The findings also established cultivar-specific differential pathogenicity of *A. solani* on tomato cultivars, with the tolerant cultivar Arka Rakshak showing minimal pathogenicity compared to the susceptible PKM-1, consistent with earlier reports (Hubballi *et al*., 2011). The morphological characteristics of *A. solani* matched the descriptions provided by the Commonwealth Mycological Institute, Kew, Surrey, England (Ellis, 1971), further substantiating the identity of the pathogen. Light microscopy also confirmed the identification of the pathogen as a member of the genus *Alternaria* (Roy *et al*., 2019).

Both susceptible and tolerant cultivars of tomato leaves pretreated with laminarin at various concentrations were evaluated for early blight severity under greenhouse conditions. The results showed a time- and dose-dependent decrease in disease severity at all tested laminarin concentrations, with the highest concentration (0.1%) showing significant reduction in disease symptoms, particularly in the tolerant cultivar. These results align with prior studies demonstrating the effectiveness of sodium alginate and methyl jasmonate methyl ester (MJME) in preventing *A. solani* infection in tomatoes (Kamakshi *et al*., 2023). Ray *et al*. (2015) documented the use of *A. solani* to screen for differences in infection patterns among tomato cultivars, highlighting the importance of basal resistance in plant-pathogen interactions (Yao *et al*., 2011). The differential responses of tolerant and susceptible cultivars observed in the present study underline their potential role in the development of disease-tolerant crops. Light microscopy analysis of tomato leaves inoculated with *A. solani* confirmed fungal infection through stomatal openings, consistent with earlier observations in *C. arietinum* (Sarkar *et al*., 2017; Mandal *et al*., 2018). The current study further documented a delay in hyphal progression in laminarin-pretreated leaves of both cultivars up to day 6, with reduced hyphal growth and penetration in the tolerant cultivar by day 8. These findings are in agreement with histopathological studies on *A. solani*-potato interactions, which have shown that infection dynamics vary based on leaf age and resistance levels (Dita *et al*., 2007; Araújo and Matsuoka, 2004).

Fluorescence microscopy further confirmed that fungal penetration was delayed in laminarin-pretreated leaves, particularly in the tolerant cultivar. By day 8, the tolerant cultivar exhibited significantly reduced fungal colonization, whereas the susceptible cultivar still showed moderate infection. Reduced hyphal development and penetration have been previously documented in tolerant crops infected with necrotrophic fungi, such as *C. arietinum* and *Botrytis cinerea* (Govrin and Levine, 2000; Ortega *et al*., 2005). This study corroborates these findings by demonstrating that laminarin treatment effectively suppresses fungal colonization in *A. solani*-infected tomato leaves, even in susceptible cultivars. The role of reactive oxygen species (ROS) in plant defense is well-documented, with ROS acting as critical signaling molecules during pathogen infection. In the current study, laminarin treatment enhanced ROS accumulation, including hydrogen peroxide (H□O□) and superoxide anion (O□□), in both susceptible and tolerant cultivars. The maximal accumulation of H□O□ in laminarin-pretreated leaves highlights its role in maintaining homeostasis and activating defense mechanisms. Elicitors such as laminarin are known to activate antioxidative systems, neutralizing the detrimental effects of ROS and preserving cellular integrity (Balmer and Mauch-Mani, 2012). The antioxidative system includes key enzymes such as peroxidases (POX), polyphenol oxidase (PPO), catalase (CAT), and superoxide dismutase (SOD), which serve as the first line of defense against ROS. The observed induction of these enzymes in laminarin-pretreated plants aligns with previous studies demonstrating their role in pathogen resistance (Mani *et al*., 2021).

Guaiacol peroxidase (GPX), in particular, plays a significant role in lignin biosynthesis, suberin deposition, and phytoalexin production, contributing to the structural fortification of plant tissues during pathogen attack. In this study, laminarin treatment induced GPX activity, with the highest levels observed in plants pretreated with 0.1% laminarin. Similar observations have been made in potatoes infected with *A. solani*, where increased GPX activity correlated with enhanced resistance (Shahbazi *et al*., 2010). Likewise, higher GPX activity in wheat cultivars resistant to *Fusarium* species underscores its critical role in limiting pathogen spread (Pieczul *et al*., 2020). Superoxide dismutase (SOD) activity also increased significantly in laminarin-treated tomato leaves. SOD is essential for dismutating superoxide radicals into H□O□ and oxygen, thereby mitigating oxidative damage. In this study, laminarin-induced SOD activity was evident in both cultivars, with a pronounced response in the tolerant cultivar. These findings are consistent with previous reports on increased SOD activity in response to pathogen infection in *Pyrus communis* (Azarabadi *et al*., 2017). Catalase (CAT) activity, which is pivotal in scavenging H□O□, was also significantly elevated in laminarin-treated plants. This enzymatic response aligns with reports that sodium alginate induces systemic acquired resistance (SAR) and enhances CAT activity during biotic stress (Dey *et al*., 2019). Enhanced CAT activity has been linked to improved pathogen resistance in several crops, including palm and tomato, further supporting its role in laminarin-induced defense (Khompatara *et al*., 2019; Elansary *et al*., 2017).

Phenylalanine ammonia-lyase (PAL) activity, a key enzyme in secondary metabolite biosynthesis, was significantly induced in laminarin-treated plants. PAL is critical for the production of lignin, flavonoids, and phenolic compounds, which contribute to cell wall reinforcement and pathogen defense. The observed increase in PAL activity is consistent with earlier findings in barley and tobacco, where PAL played a pivotal role in resistance to fungal pathogens (Mishra *et al*., 2011; Kaur *et al*., 2021). Polyphenol oxidase (PPO) activity also showed a significant increase in laminarin-treated plants. PPO-mediated production of quinones disrupts pathogen signaling and enhances defense responses. These findings align with studies in pearl millet and other crops, where PPO activity was associated with resistance to fungal infections (Raj *et al*., 2006; Agostinetto *et al*., 2016).

Altogether, this study explored the efficacy of laminarin, an algal-derived polysaccharide, in inducing defense responses against early blight disease caused by *A*.*solani* in tomato cultivars. The investigation involved both susceptible (PKM-1) and tolerant (Arka Rakshak) cultivars, with detailed analyses of disease progression, ROS accumulation, histopathological changes, and antioxidant enzyme activities. Pathogenicity tests confirmed the virulence of *A. solani*, and morphological traits of the pathogen were consistent with established descriptions. Disease severity was significantly reduced in laminarin-pretreated plants, with a dose- and time-dependent effect, particularly at a 0.1% concentration. This reduction was more pronounced in the tolerant cultivar, which exhibited minimal disease symptoms and delayed fungal colonization. Histopathological studies revealed suppressed fungal growth, limited hyphal penetration, and delayed symptom progression in laminarin-treated plants. Fluorescence microscopy corroborated these findings, showing reduced fungal colonization in laminarin-pretreated leaves, especially in the tolerant cultivar. Laminarin also induced a robust oxidative burst, characterized by the accumulation of hydrogen peroxide (H□O□) and superoxide anion (O□·□), which are known to play critical roles in plant defense signaling. Antioxidative enzymes, including GPX, PPO, SOD, CAT, and PAL, were significantly activated in laminarin-treated plants, underscoring the enhanced defense response.

These results demonstrate that laminarin effectively reduces early blight disease symptoms by enhancing innate resistance mechanisms in tomato plants. It induces ROS-mediated defense pathways, activates antioxidative enzymes, and reinforces structural and biochemical barriers against *A. solani*. These findings align with previous research on algal polysaccharides and highlight laminarin’s potential as an eco-friendly and sustainable alternative to synthetic fungicides. In conclusion, laminarin serves as a potent elicitor of defense mechanisms in tomato plants, offering a promising strategy for managing early blight disease while promoting sustainable agricultural practices. Its application could reduce reliance on chemical fungicides, thereby contributing to environmental and food safety.

## Supporting information

Total Supplementary Data

## ACKNOWLEDGEMENTS

The authors wish to thank The Director, Centre for Advanced Studies in Botany, University of Madras for providing the laboratory facility to carry out the experiments. The corresponding author also thanks the Tamil Nadu State Council for Higher Education (TANSCHE), Chennai, Tamil Nadu in India for providing financial support.

## REFERENCES

1. Agostinetto, D., Perboni, L.T., Langaro, A.C., Gomes, J., Fraga, D.S. and Franco, J.J., 2016. Changes in photosynthesis and oxidative stress in wheat plants submmited to herbicides application. PlantaDaninha, 34, pp.01–09.

2. Ali, O., Ramsubhag, A. and Jayaraman, J., 2021. Biostimulant properties of seaweed extracts in plants: Implications towards sustainable crop production. Plants, 10(3), p.531.

3. Araujo, J.C.A.D. and Matsuoka, K., 2004. Histopathology of the interaction between Alternaria solani and resistant and susceptible tomato plants. Fitopatologia Brasileira, 29, pp.268–275.

4. Arnnok, P., Ruangviriyachai, C., Mahachai, R., Techawongstien, S. and Chanthai, S., 2010. Optimization and determination of polyphenol oxidase and peroxidase activities in hot pepper (Capsicum annuum L.) pericarb. Int. food res. J, 17, pp.385–392.

5. Azarabadi, S., Abdollahi, H., Torabi, M., Salehi, Z. and Nasiri, J., 2017. ROS generation, oxidative burst and dynamic expression profiles of ROS-scavenging enzymes of superoxide dismutase (SOD), catalase (CAT) and ascorbate peroxidase (APX) in response to Erwinia amylovora in pear (Pyrus communis L). European Journal of Plant Pathology, 147, pp.279–294.

6. Balmer, D. and Mauch-Mani, B., 2012. Plant hormones and metabolites as universal vocabulary in plant defense signaling. Biocommunication of Plants, pp.37–50.

7. Beauchamp, C. and Fridovich, I., 1971. Superoxide dismutase: improved assays and an assay applicable to acrylamide gels. Analytical biochemistry, 44(1), pp.276–287.

8. Bhattacharjee, C., Saxena, V.K. and Dutta, S., 2017. Fruit juice processing using membrane technology: A review. Innovative Food Science & Emerging Technologies, 43, pp.136–153.

9. Buchanan□Wollaston, V., Earl, S., Harrison, E., Mathas, E., Navabpour, S., Page, T. and Pink, D., 2003. The molecular analysis of leaf senescence–a genomics approach. Plant biotechnology journal, 1(1), pp.3–22.

10. Chen, Z., Silva, H. and Klessig, D.F., 1993. Active oxygen species in the induction of plant systemic acquired resistance by salicylic acid. Science, 262(5141), pp.1883–1886.

11. Das, K. and Roychoudhury, A., 2014. Reactive oxygen species (ROS) and response of antioxidants as ROS-scavengers during environmental stress in plants. Frontiers in environmental science, 2, p.53.

12. Daudi, A. and O’Brien, J.A., 2012. Detection of hydrogen peroxide by DAB staining in Arabidopsis leaves. Bio-protocol, 2(18), pp. e263–e263.

13. Dey, P., Ramanujam, R., Venkatesan, G. and Nagarathnam, R., 2019. Sodium alginate potentiates antioxidant defense and PR proteins against early blight disease caused by Alternaria solani in Solanum lycopersicum Linn. PLoS One, 14(9), p.e0223216.

14. Di, X., Gomila, J.O. and Takken, F.L., 2017. Involvement of salicylic acid, ethylene and jasmonic acid signalling pathways in the susceptibility of tomato to Fusarium oxysporum. Molecular Plant Pathology, 18(7), pp.1024–1035.

15. Dita, M.A., Brommonschenkel, S.H., Matsuoka, K. and Mizubuti, E.S.G., 2007. Histopathological study of the Alternaria solani infection process in potato cultivars with different levels of early blight resistance. Journal of Phytopathology, 155(7□8), pp.462–469.

16. Elansary, H.O., Yessoufou, K., Abdel-Hamid, A.M., El-Esawi, M.A., Ali, H.M. and Elshikh, M.S., 2017. Seaweed extracts enhance salam turfgrass performance during prolonged irrigation intervals and saline shock. Frontiers in Plant Science, 8, p.830.

17. Ellis, M. B., & Hyphomycetes, D. (1971). Commonwealth Mycological Institute. Kew, Surrey, England.

18. Gao, Z., Zhang, B., Liu, H., Han, J., & Zhang, Y. (2017). Identification of endophytic Bacillus velezensis ZSY-1 strain and antifungal activity of its volatile compounds against Alternaria solani and Botrytis cinerea. Biological Control, 105, 27–39.

19. Gauthier, A., Trouvelot, S., Kelloniemi, J., Frettinger, P., Wendehenne, D., Daire, X., Joubert, J.M., Ferrarini, A., Delledonne, M., Flors, V. and Poinssot, B., 2014. The sulfated laminarin triggers a stress transcriptome before priming the SA-and ROS-dependent defenses during grapevine’s induced resistance against Plasmopara viticola. PLoS One, 9(2), p.e88145.

20. Goel, N. and Paul, P.K., 2015. Polyphenol oxidase and lysozyme mediate induction of systemic resistance in tomato, when a bioelicitor is used. Journal of Plant Protection Research, 55(4).

21. Govrin, E. M., & Levine, A. (2000). The hypersensitive response facilitates plant infection by the necrotrophic pathogen Botrytis cinerea. Current biology, 10(13), 751–757.

22. Hu, X., Wansha, L., Chen, Q. and Yang, Y., 2009. Early signals transduction linking the synthesis of jasmonic acid in plant. Plant Signaling and Behavior, 4(8), pp.696–697.

23. Hubballi, M., Nakkeeran, S., Raguchander, T., Anand, T., & Renukadevi, P. (2011). Physiological characterisation of Colletotrichum gloeosporioides, the incitant of anthracnose disease of noni in India. Archives of phytopathology and plant protection, 44(11), 1105–1114.

24. Irulappan, V., Kandpal, M., Saini, K., Rai, A., Ranjan, A., Sinharoy, S. and Senthil-Kumar, M., 2022. Drought stress exacerbates fungal colonization and endodermal invasion and dampens defense responses to increase dry root rot in chickpea. Molecular Plant-Microbe Interactions, 35(7), pp.583–591.

25. Javed, T., Shabbir, R., Ali, A., Afzal, I., Zaheer, U., & Gao, S. J. (2020). Transcription factors in plant stress responses: Challenges and potential for sugarcane improvement. Plants, 9(4), 491.

26. Jendresen, C. B., Stahlhut, S. G., Li, M., Gaspar, P., Siedler, S., Förster, J., … & Nielsen, A. T. (2015). Highly active and specific tyrosine ammonia-lyases from diverse origins enable enhanced production of aromatic compounds in bacteria and Saccharomyces cerevisiae. Applied and environmental microbiology, 81(13), 4458–4476.

27. Jones, J.D. and Dangl, J.L., 2006. The plant immune system. nature, 444(7117), pp.323–329.

28. Kamakshi, K., MohanaPrasad, J., Muthamilarasan, M., & Radhakrishnan, N. (2023). Foliar application of Methyl Jasmonate Methyl Ester elicits differential antioxidant defence and expression of defence□related genes against early blight disease of tomato. Journal of Phytopathology.

29. Kaur, A., Sharma, V. K., & Sharma, S. (2021). Management of spot blotch of barley: an eco-friendly approach. Australasian Plant Pathology, 50, 389–401.

30. Khan, K. A., Shoaib, A., ArshadAwan, Z., Basit, A., & Hussain, M. (2018). Macrophominaphaseolina alters the biochemical pathway in Vignaradiata chastened by Zn2+ and FYM to improve plant growth. Journal of Plant Interactions, 13(1), 131–140.

31. Khompatara, K., Pettongkhao, S., Kuyyogsuy, A., Deenamo, N., & Churngchow, N. (2019). Enhanced resistance to leaf fall disease caused by Phytophthorapalmivora in rubber tree seedling by Sargassumpolycystum extract. Plants, 8(6), 168.

32. Koley, S., Mahapatra, S.S. and Kole, P.C., 2015. In vitro efficacy of bio-control agents and botanicals on the growth inhibition of Alternaria solani causing early leaf blight of tomato. International Journal of Bio-resource, Environment and Agricultural Sciences, 1(3), pp.114–118.

33. Kumar, D., Yusuf, M. A., Singh, P., Sardar, M., & Sarin, N. B. (2014). Histochemical detection of superoxide and H2O2 accumulation in Brassica juncea seedlings. Bioprotocol, 4(8), e1108–e1108.

34. Lavania, M., Chauhan, P.S., Chauhan, S.V.S., Singh, H.B. and Nautiyal, C.S., 2006. Induction of plant defense enzymes and phenolics by treatment with plant growth– promoting rhizobacteria Serratia marcescens NBRI1213. Current Microbiology, 52(5), pp.363–368.

35. Lavanya, S. N., Niranjan-Raj, S., Jadimurthy, R., Sudarsan, S., Srivastava, R., Tarasatyavati, C., … & Nayaka, S. C. (2022). Immunity elicitors for induced resistance against the downy mildew pathogen in pearl millet. Scientific Reports, 12(1), 4078.

36. Li, X., Wang, Y., Chen, S., Tian, H., Fu, D., Zhu, B., … & Zhu, H. (2018). Lycopene is enriched in tomato fruit by CRISPR/Cas9-mediated multiplex genome editing. Frontiers in plant science, 9, 559.

37. Mandal, S., Rajarammohan, S., & Kaur, J. (2018). Alternaria brassicae interactions with the model Brassicaceae member Arabidopsis thaliana closely resembles those with Mustard (Brassica juncea). Physiology and Molecular Biology of Plants, 24, 51–59.

38. Mani, S. D., & Nagarathnam, R. (2018). Sulfated polysaccharide from Kappaphycus alvarezii (Doty) Doty ex PC Silva primes defense responses against anthracnose disease of Capsicum annuum Linn. Algal research, 32, 121–130.

39. Mani, S. D., Govindan, M., Muthamilarasan, M., & Nagarathnam, R. (2021). A sulfated polysaccharide κ-carrageenan induced antioxidant defense and proteomic changes in chloroplast against leaf spot disease of tomato. Journal of Applied Phycology, 33, 2667–2681.

40. Mani, S. D., Pandey, S., Govindan, M., Muthamilarasan, M., & Nagarathnam, R. (2021). Transcriptome dynamics underlying elicitor-induced defense responses against Septoria leaf spot disease of tomato (Solanum lycopersicum L.). Physiology and Molecular Biology of Plants, 27, 873–888.

41. Mishra, V. K., Biswas, S. K., & Rajik, M. (2011). Biochemical mechanism of resistance to alternaria blight by different varieties of wheat. International Journal of Plant Pathology, 2(2), 72–80.

42. Mittler, R., Vanderauwera, S., Gollery, M. and Van Breusegem, F., 2004. Reactive oxygen gene network of plants. Trends in Plant Science, 9(10), pp.490–498.

43. Mugao, L. (2023). Morphological and Molecular Variability of Alternaria solani and Phytophthora infestans Causing Tomato Blights. International Journal of Microbiology, 2023.

44. Ortega, X., Velásquez, J. C., & Pérez, L. M. (2005). IP3 production in the hypersensitive response of lemon seedlings against Alternariaalternata involves active protein tyrosine kinases but not a G-protein. Biological Research, 38(1), 89–99.

45. Pieczul, K., Dobrzycka, A., Wolko, J., Perek, A., Zielezińska, M., Bocianowski, J., & Rybus-Zając, M. (2020). The activity of β-glucosidase and guaiacol peroxidase in different genotypes of winter oilseed rape (Brassica napus L.) infected by Alternaria black spot fungi. ActaPhysiologiaePlantarum, 42, 1–9.

46. Radhakrishnan, N. and Balasubramanian, R., 2009. Salicylic acid induced defence responses in Curcuma longa (L.) against Pythium aphanidermatum infection. Crop Protection, 28(11), pp.974–979.

47. Raj, S. N., Sarosh, B. R., & Shetty, H. S. (2006). Induction and accumulation of polyphenol oxidase activities as implicated in development of resistance against pearl millet downy mildew disease. Functional Plant Biology, 33(6), 563–571.

48. Rashmi, T. and Vishunavat, K., 2012. Management of early blight (Alternaria solani) in tomato by integration of fungicides and cultural practices. International Journal of Plant Protection, 5(2), pp.201–206.

49. Ray, S., Mondal, S., Chowdhury, S. and Kundu, S., 2015. Differential responses of resistant and susceptible tomato varieties to inoculation with Alternaria solani. Physiological and Molecular Plant Pathology, 90, pp.78–88.

50. Roy, C. K., Akter, N., Sarkar, M. K., Pk, M. U., Begum, N., Zenat, E. A., & Jahan, M. A. (2019). Control of early blight of tomato caused by and screening of tomato varieties against the pathogen. The Open Microbiology Journal, 13(1).

51. Sak, M., Dokupilová, I., Kaňuková, Š., Mrkvová, M., Mihálik, D., Hauptvogel, P., & Kraic, J. (2021). Biotic and abiotic elicitors of stilbenes production in Vitisvinifera L. cell culture. Plants, 10(3), 490.

52. Sarkar, D., Maji, R. K., Dey, S., Sarkar, A., Ghosh, Z., & Kundu, P. (2017). Integrated miRNA and mRNA expression profiling reveals the response regulators of a susceptible tomato cultivar to early blight disease. DNA Research, 24(3), 235–250.

53. Shah, J., Chaturvedi, R., Chowdhury, Z., Venables, B. and Petros, R.A., 2014. Signaling by small metabolites in systemic acquired resistance. The Plant Journal, 79(4), pp.645–658.

54. Shahbazi, H., Aminian, H., Sahebani, N. and Halterman, D.A., 2010. Biochemical evaluation of resistance responses of potato to different isolates of Alternaria solani. Phytopathology, 100(5), pp.454–459.

55. Shetty, H. S., Suryanarayan, S. M., Jogaiah, S., Janakirama, A. R. S., Hansen, M., Jørgensen, H. J. L., & Tran, L. S. P. (2019). Bioimaging structural signatures of the oomycete pathogen Sclerosporagraminicola in pearl millet using different microscopic techniques. Scientific reports, 9(1), 15175.

56. Shukla, P. S., Borza, T., Critchley, A. T., & Prithiviraj, B. (2021). Seaweed-based compounds and products for sustainable protection against plant pathogens. Marine drugs, 19(2), 59.

57. Sibounnavong, P., Keoudone, C., Soytong, K., Divina, C. C., & Kalaw, S. P. (2010). A new mycofungicide from Emericellanidulans against tomato wilt caused by Fusarium oxysporum f. sp. lycopersici in vivo. Journal of Agricultural Technology, 6(1), 19–30.

58. Thinakaran, A. M., Adaikkalam, C., Asaithambi, V., & Nagarathinam, R. (2020). Sulfamethoxazole activates antioxidant defense response against Alternaria solani infection in Solanum lycopersicum Linn. Archives of Phytopathology and Plant Protection, 53(17-18), 806–827.

59. Tziros, G. T., Samaras, A., & Karaoglanidis, G. S. (2021). Laminarin induces defense responses and efficiently controls olive leaf spot disease in olive. Molecules, 26(4), 1043.

60. Volk, S. and Feierabend, J., 1989. Photoinactivation of catalase at low temperature and its relevance to photosynthetic and peroxide metabolism in leaves. Plant, Cell and Environment, 12(7), pp.701–712.

61. Wanke, A., Rovenich, H., Schwanke, F., Velte, S., Becker, S., Hehemann, J. H., … & Zuccaro, A. (2020). Plant species□specific recognition of long and short β□1, 3□linked glucans is mediated by different receptor systems. The Plant Journal, 102(6), 1142–1156.

62. War, A.R., Paulraj, M.G., War, M.Y. and Ignacimuthu, S., 2011. Role of salicylic acid in induction of plant defense system in chickpea (Cicer arietinum L.). Plant Signaling and Behavior, 6(11), pp.1787–1792.

63. Wiesel, L., Newton, A. C., Elliott, I., Booty, D., Gilroy, E. M., Birch, P. R., & Hein, I. (2014). Molecular effects of resistance elicitors from biological origin and their potential for crop protection. Frontiers in plant science, 5, 655.

64. Wu, Y.R., Lin, Y.C. and Chuang, H.W., 2016. Laminarin modulates the chloroplast antioxidant system to enhance abiotic stress tolerance partially through the regulation of the defensin-like gene expression. Plant Science, 247, pp.83-92.

65. Xin, Z., Cai, X., Chen, S., Luo, Z., Bian, L., Li, Z., Ge, L. and Chen, Z., 2019. A disease resistance elicitor laminarin enhances tea defense against a piercing herbivore Empoasca (Matsumurasca) onukii Matsuda. Scientific Reports, 9(1), pp.1–13.

66. Yang, T., Qiu, L., Huang, W., Xu, Q., Zou, J., Peng, Q., … & Xi, D. (2020). Chilli veinal mottle virus HCPro interacts with catalase to facilitate virus infection in Nicotianatabacum. Journal of experimental botany, 71(18), 5656–5668.

67. Yao, Z., Rashid, K. Y., Adam, L. R., & Daayf, F. (2011). Verticillium dahliae’s VdNEP acts both as a plant defence elicitor and a pathogenicity factor in the interaction with Helianthus annuus. Canadian journal of plant pathology, 33(3), 375–388.

68. You, M. P., Lamichhane, J. R., Aubertot, J. N., & Barbetti, M. J. (2020). Understanding why effective fungicides against individual soilborne pathogens are ineffective with soilborne pathogen complexes. Plant disease, 104(3), 904–920.

69. Zhao, H., Sun, X., Xue, M., Zhang, X. and Li, Q., 2016. Antioxidant enzyme responses induced by whiteflies in tobacco plants in defense against aphids: catalase may play a dominant role. PLoS One, 11(10), p.e0165454.

