## Supplementary material for "A Biopolymer Laminarin Elicits Antioxidant Defense in Different Cultivars of *Solanum Lycopersicum* Against Early Blight Disease Caused by *Alternaria Solani*": Total Supplementary Data: Supplementary data.docx

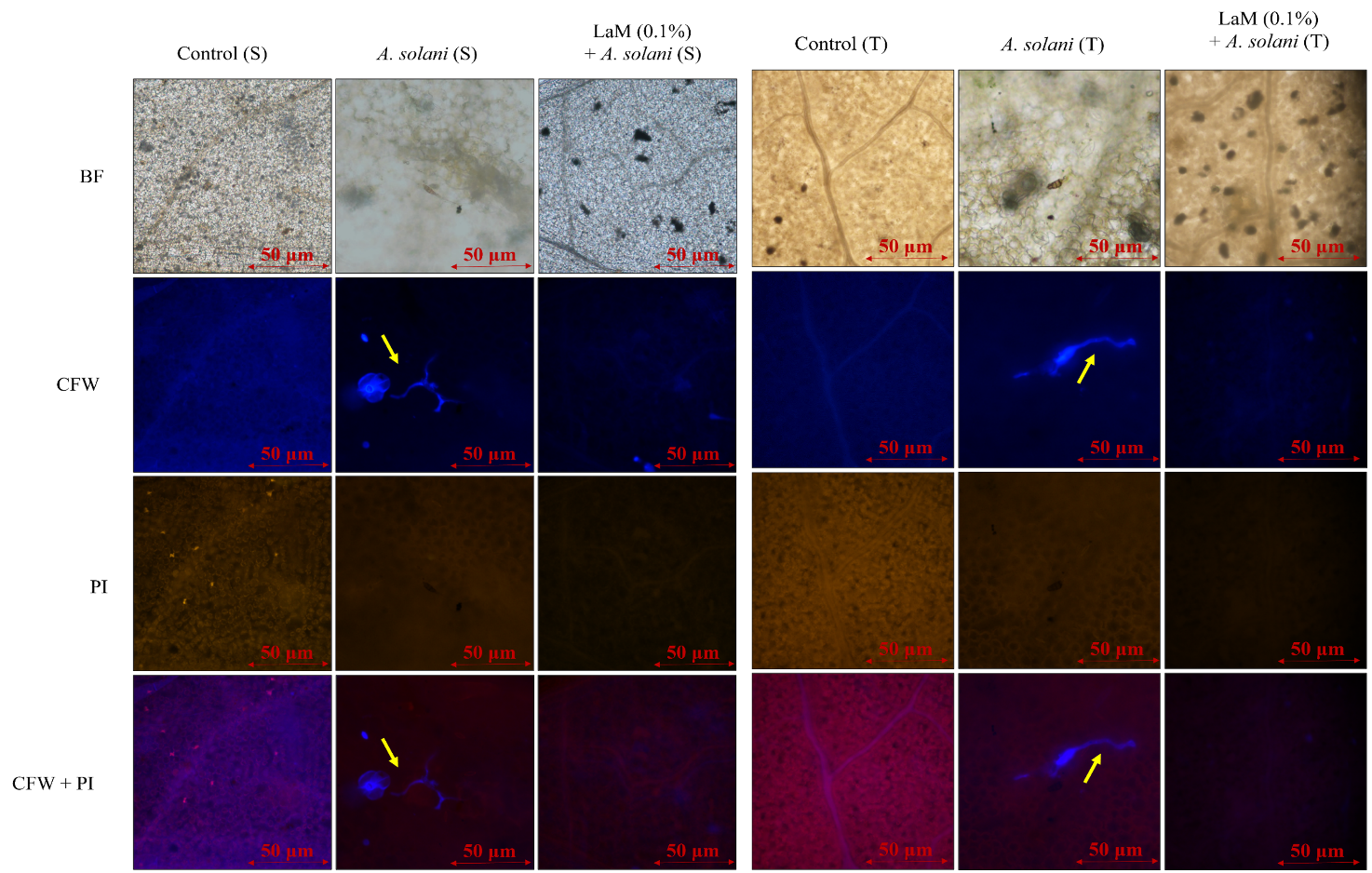
 **Fig.S-1.** Fluorescence microscopic analysis of *A. solani* infection patterns on *S. lycopersicum*
(PKM-1, susceptible cultivar) and (Arka Rakshak, tolerant cultivar) pretreated with laminarin and followed by *A. solani.* Samples collected after the first visible symptoms appeared at day 2.
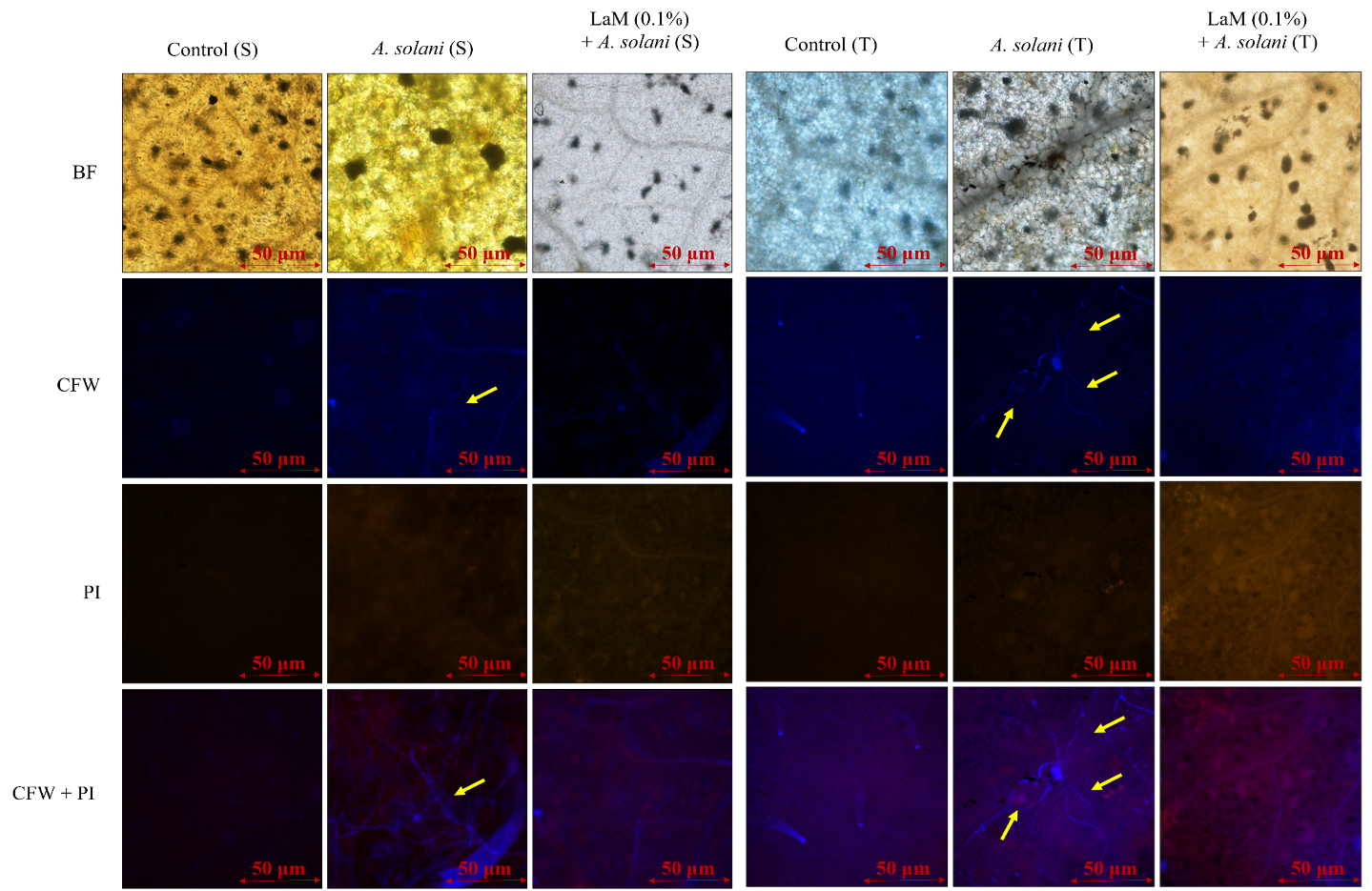

**Fig.S-2.** Fluorescence microscopic analysis of *A. solani* infection patterns on *S. lycopersicum*
(PKM-1, susceptible cultivar) and (Arka Rakshak, tolerant cultivar) pretreated with laminarin and followed by *A. solani.* Samples collected after the first visible symptoms appeared at day 4.

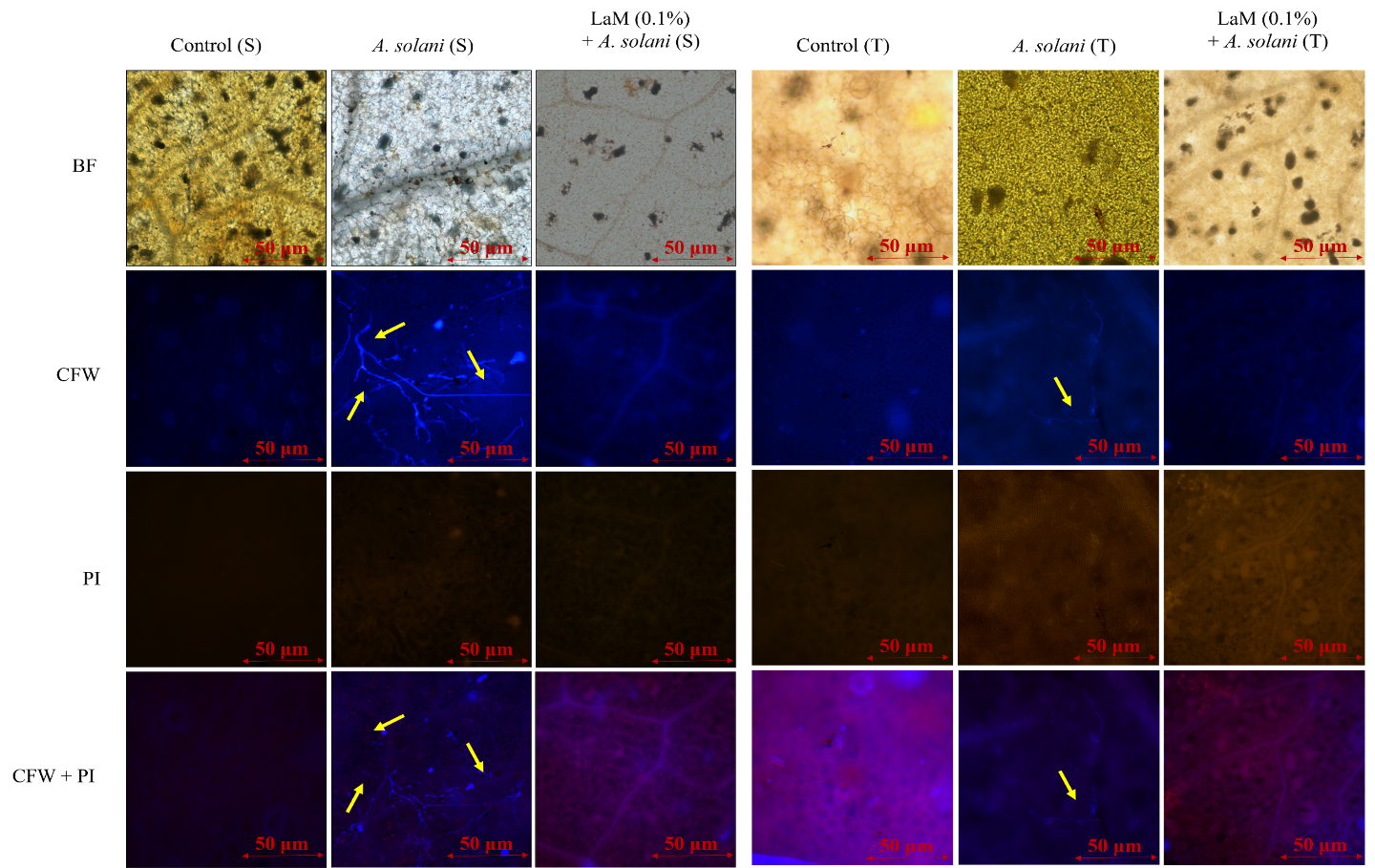

**Fig.S-3.** Fluorescence microscopic analysis of *A. solani* infection patterns on *S. lycopersicum*
(PKM-1, susceptible cultivar and Arka Rakshak, tolerant cultivar) pretreated with laminarin and followed by
*A. solani.* Samples collected after the first visible symptoms appeared at day 6.

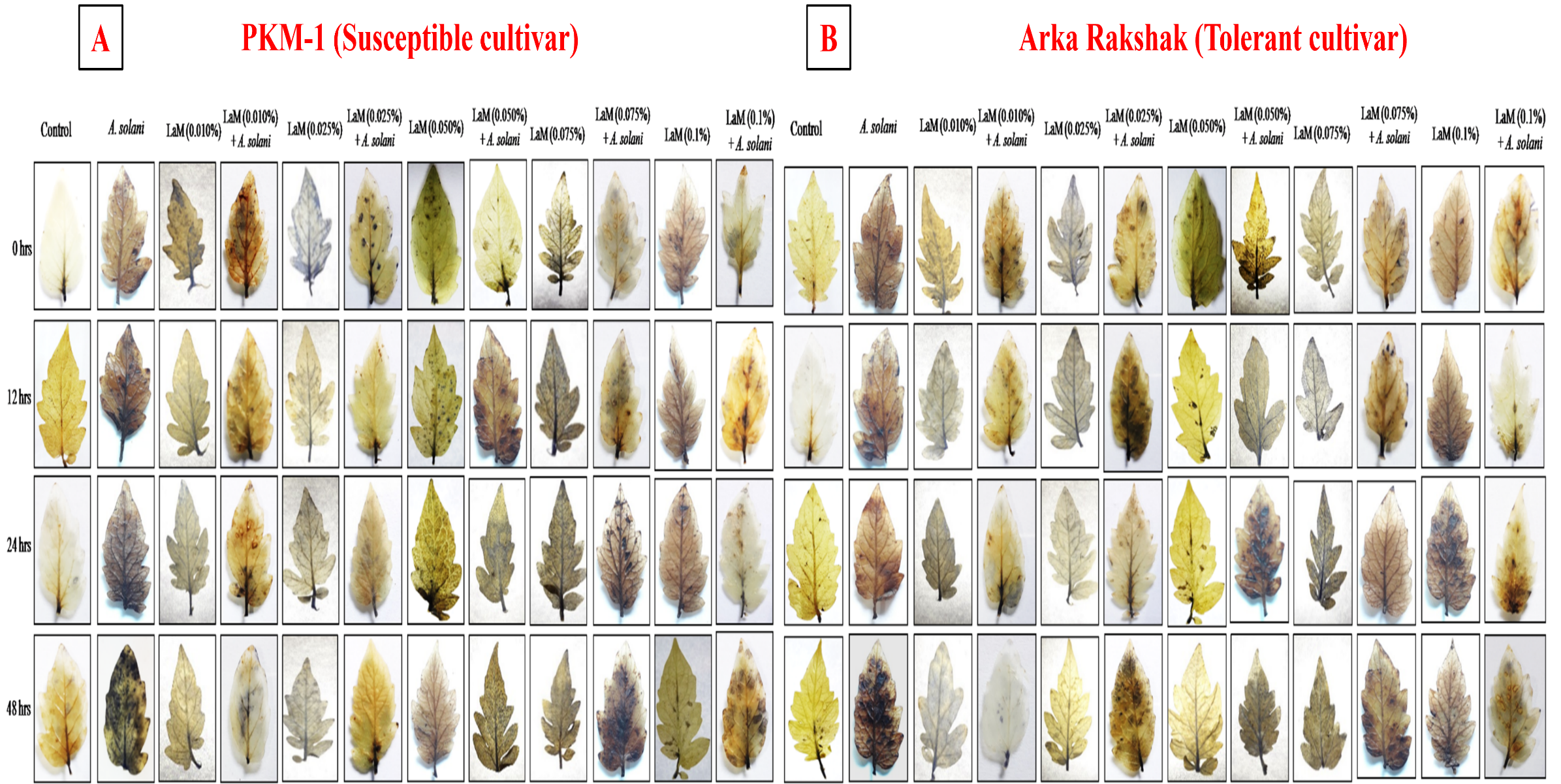
**Fig.S-4A.** *In-situ* histochemical localization of superoxide anion (O2^•−^) accumulation on *S. lycopersicum* (PKM-1, susceptible cultivar) leaves pretreated with different concentrations of laminarin and samples collected at 0 hrs, 12 hrs, 24 hrs and 48 hrs.

**Fig.S-4B.** *In-situ* histochemical localization of superoxide anion (O2^•−^) accumulation on *S. lycopersicum* (Arka Rakshak, tolerant cultivar) leaves pretreated with different concentrations of laminarin and samples collected at 0 hrs, 12 hrs, 24 hrs and 48 hrs.

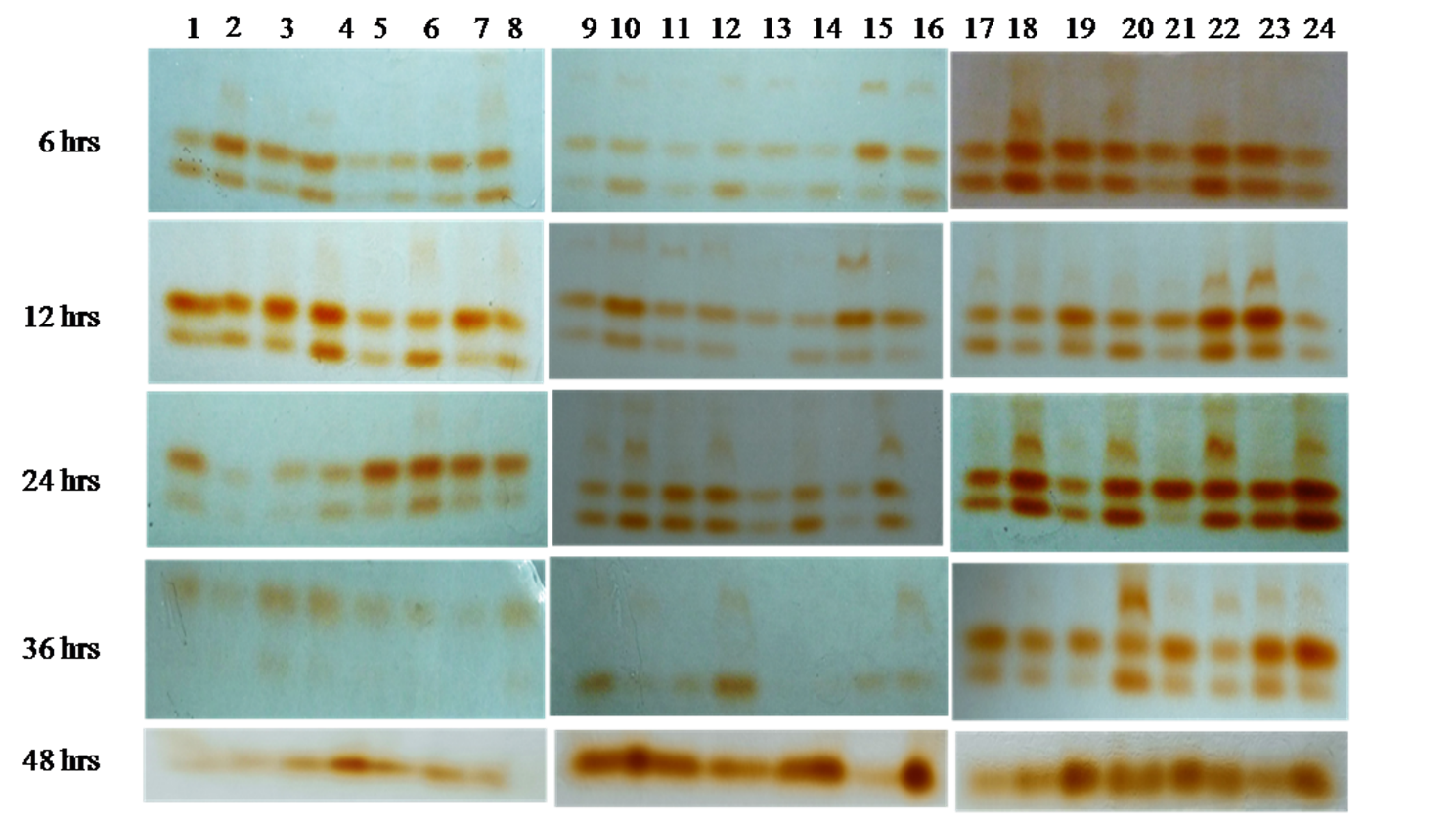

**Fig.S-5.** Native-PAGE analysis of Guaiacol peroxidase (GPX) isozymes expressions of
*S. lycopersicum* leaves pretreated with different concentrations of laminarin and infected with
*A. solani*.

1. Control (S), 2. Control (T), 3. *A. solani* infected (S),
4. *A. solani* infected (T), 5. Laminarin (0.010%) (S), 6. Laminarin (0.010%) (T),
7. Laminarin (0.010%) + *A. solani* infected (S), 8. Laminarin (0.010%) + *A. solani* infected (T), 9. Laminarin (0.025%) (S), 10. Laminarin (0.025%) (T), 11. Laminarin (0.025%) + *A. solani* infected (S), 12. Laminarin (0.025%) + *A. solani* infected (T), 13. Laminarin (0.050%) (S),
14. Laminarin (0.050%) (T), 15. Laminarin (0.050%) + *A. solani* infected (S), 16. Laminarin (0.050%) + *A. solani* infected (T), 17. Laminarin (0.075%) (S), 18. Laminarin (0.075%) (T),
19. Laminarin (0.075%) + *A. solani* infected (S), 20. Laminarin (0.075%) + *A. solani*
infected (T), 21. Laminarin (0.1%) (S), 22. Laminarin (0.1%) (T), 23. Laminarin (0.1%) +
*A. solani* infected (S), 24. Laminarin (0.1%) + *A. solani* infected (T). S - susceptible cultivar and
T - tolerant cultivar.

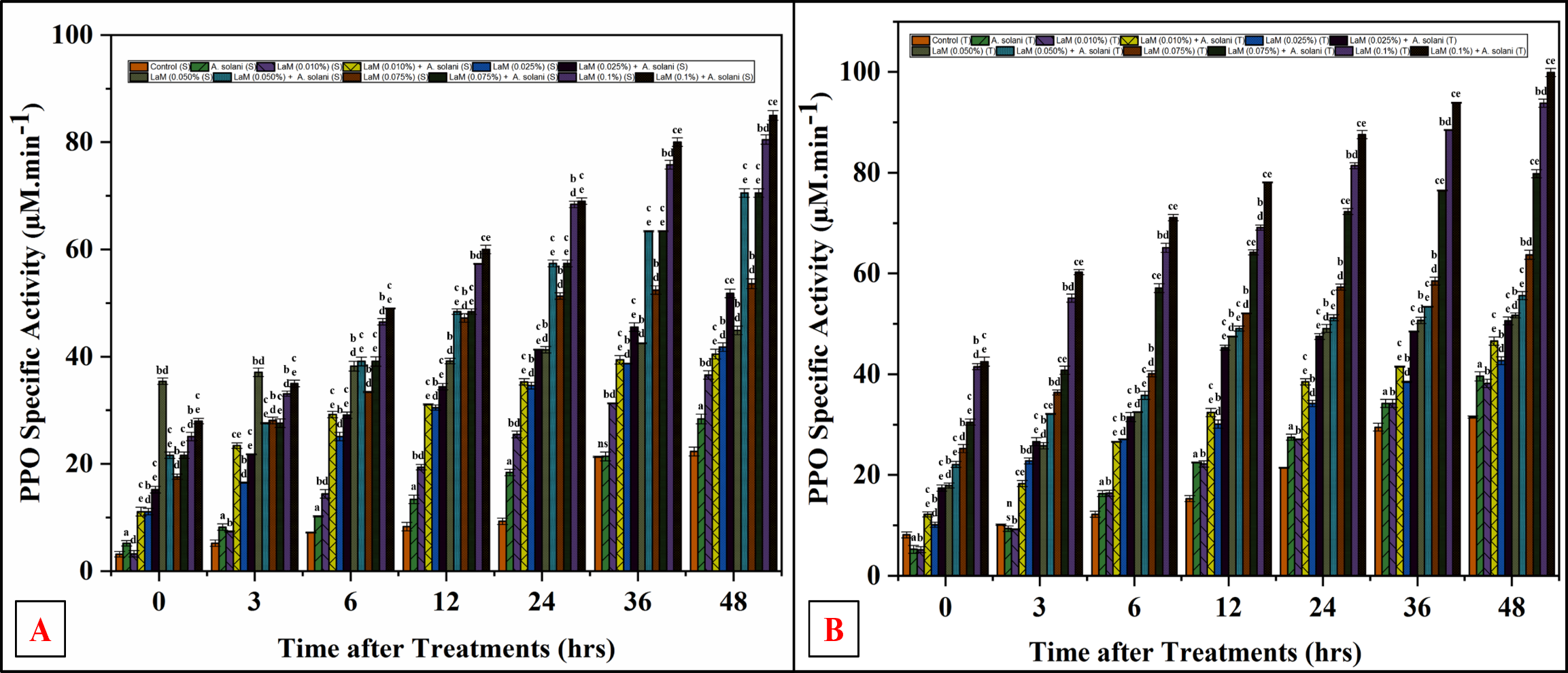
**Fig.S-6A.** Polyphenol oxidase (PPO) activity of *S. lycopersicum* (PKM-1, susceptible cultivar) leaves pretreated with different concentrations of laminarin and infected with *A. solani.*

**Fig.S-6B.** Polyphenol oxidase (PPO) activity of *S. lycopersicum* (Arka Rakshak, tolerant cultivar) leaves pretreated with different concentrations of laminarin and infected with *A. solani.*

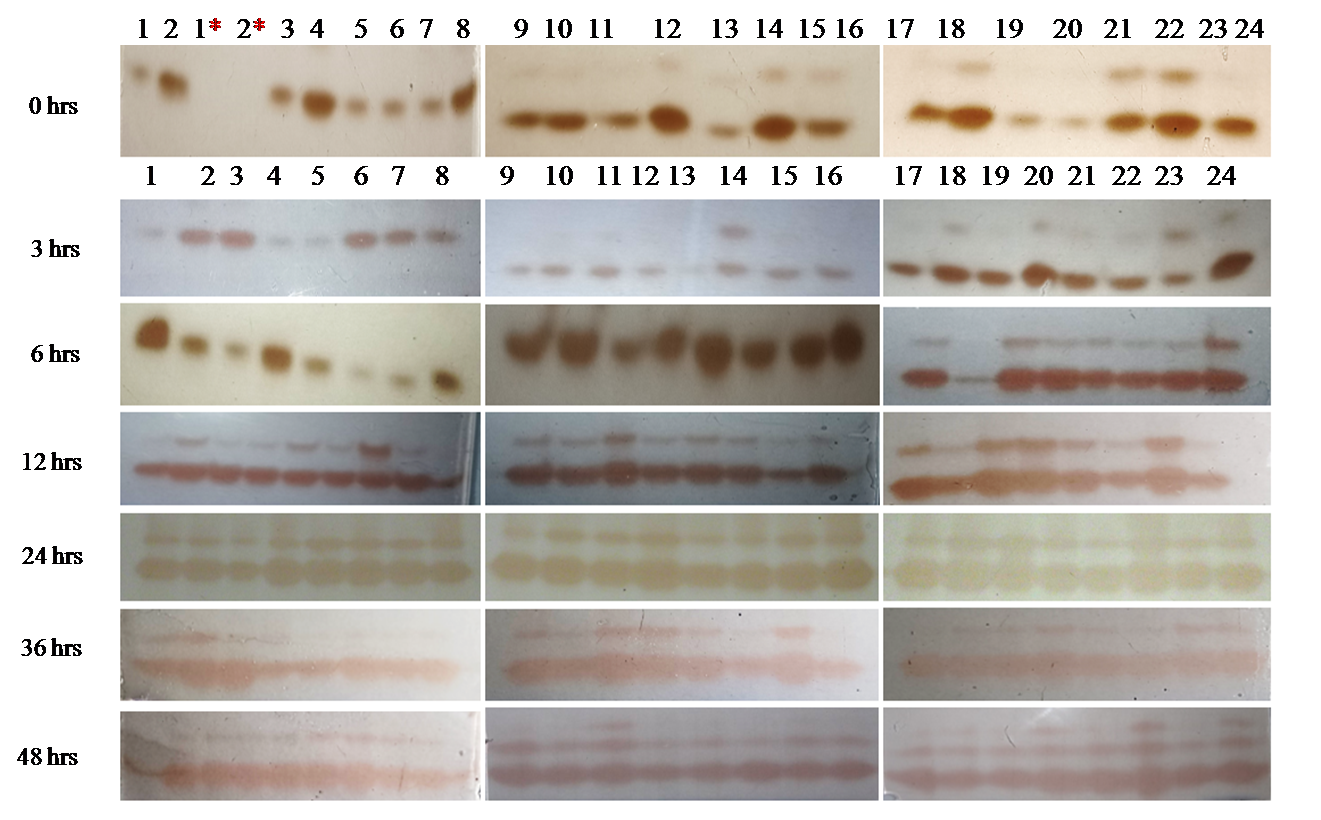
**Fig.S-7.** Native-PAGE analysis of Polyphenol oxidase (PPO) isozymes expressions of
*S. lycopersicum* leaves pretreated with different concentrations of laminarin and infected with
*A. solani.*

1. Control (S), 2. Control (T), 1*. Control (S) (PPO inhibitor, Citric acid (5%)), 2*. Control (T)
(PPO inhibitor, Citric acid (5%)), 3. *A. solani* infected (S), 4. *A. solani* infected (T),
5. Laminarin (0.010%) (S), 6. Laminarin (0.010%) (T), 7. Laminarin (0.010%) +
*A. solani* infected (S), 8. Laminarin (0.010%) + *A. solani* infected (T), 9. Laminarin (0.025%) (S), 10. Laminarin (0.025%) (T), 11. Laminarin (0.025%) + *A. solani* infected (S),
12. Laminarin (0.025%) + *A. solani* infected (T), 13. Laminarin (0.050%) (S),
14. Laminarin (0.050%) (T), 15. Laminarin (0.050%) + *A. solani* infected (S),
16. Laminarin (0.050%) + *A. solani* infected (T), 17. Laminarin (0.075%) (S),
18. Laminarin (0.070%) (T), 19. Laminarin (0.075%) + *A. solani* infected (S),
20. Laminarin (0.075%) + *A. solani* infected (T), 21. Laminarin (0.1%) (S), 22. Laminarin (0.1%) (T), 23. Laminarin (0.1%) + *A. solani* infected (S), 24. Laminarin (0.1%) + *A. solani*
infected (T). S - susceptible cultivar and T - tolerant cultivar.

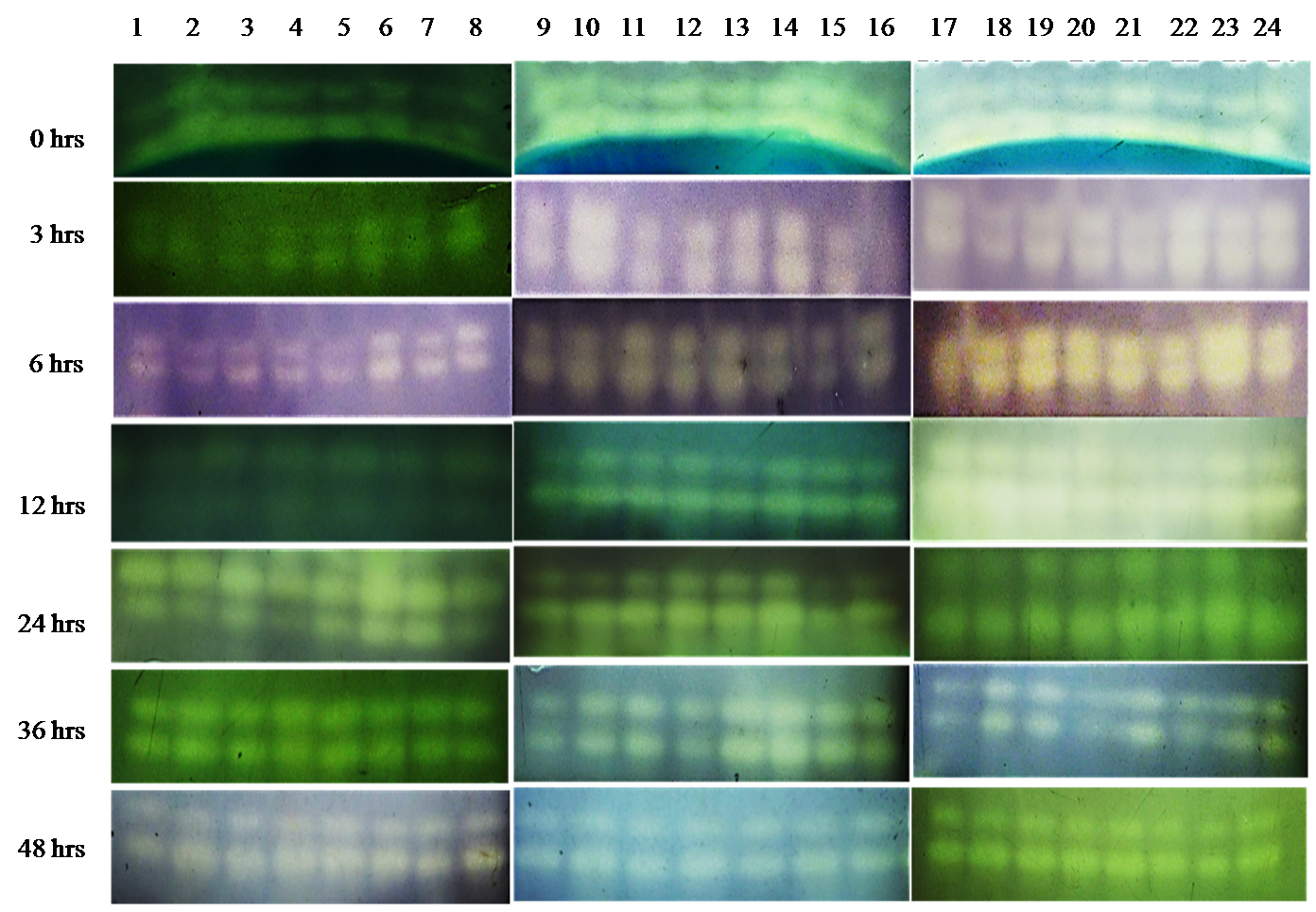
**Fig.S-8.** Native-PAGE analysis of Superoxide Dismutase (SOD) isoenzymes expressions of
*S. lycopersicum* leaves pretreated with different concentrations of laminarin and infected with
*A. solani.*

1. Control (S), 2. Control (T), 3. A. solani infected (S), 4. A. solani infected (T),
5. Laminarin (0.010%) (S), 6. Laminarin (0.010%) (T), 7. Laminarin (0.010%) + *A. solani* infected (S), 8. Laminarin (0.010%) + A. solani infected (T), 9. Laminarin (0.025%) (S),
10. Laminarin (0.025%) (T), 11. Laminarin (0.025%) + *A. solani* infected (S), 12. Laminarin (0.025%) + *A. solani* infected (T), 13. Laminarin (0.050%) (S), 14. Laminarin (0.050%) (T),
15. Laminarin (0.050%) + *A. solani* infected (S), 16. Laminarin (0.050%) + *A. solani* infected (T), 17. Laminarin (0.075%) (S), 18. Laminarin (0.070%) (T), 19. Laminarin (0.075%) +
*A. solani* infected (S), 20. Laminarin (0.075%) + *A. solani* infected (T), 21. Laminarin (0.1%) (S), 22. Laminarin (0.1%) (T), 23. Laminarin (0.1%) + *A. solani* infected (S),
24. Laminarin (0.1%) + *A. solani* infected (T). S - susceptible cultivar and T - tolerant cultivar.

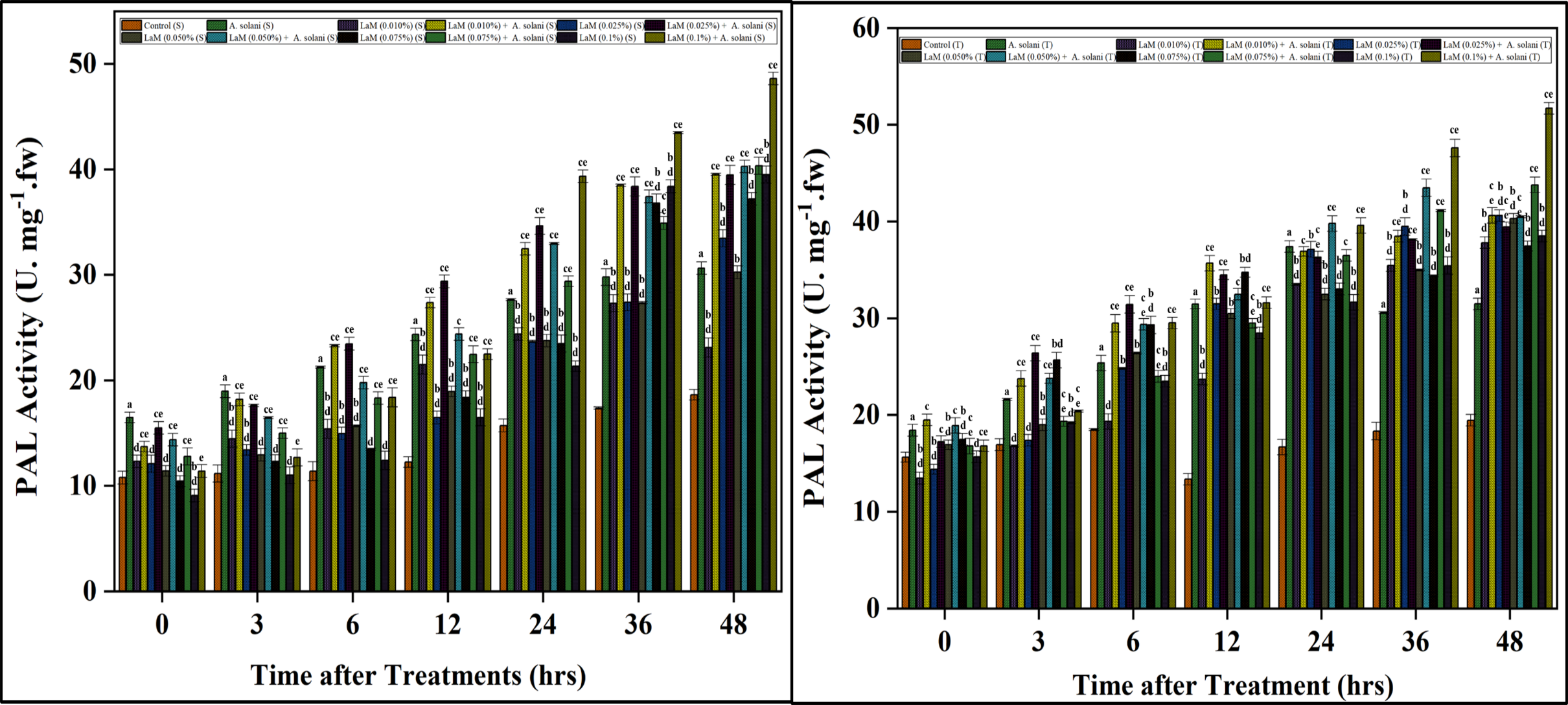

**Fig.S-9A.** Phenylalanine ammonia-lyase activity (PAL) activity of *S. lycopersicum* (PKM-1, susceptible cultivar) leaves pretreated with different concentrations of laminarin and infected with *A. solani.*

**Fig.S-9B.** Phenylalanine ammonia-lyase activity (PAL) activity of *S. lycopersicum* (Arka Rakshak, tolerant cultivar) leaves pretreated with different concentrations of laminarin and infected with *A. solani.*
